# Multi-layered dosage compensation of the avian Z chromosome

**DOI:** 10.1101/2024.08.20.608780

**Authors:** Natali Papanicolaou, Antonio Lentini, Sebastian Wettersten, Michael Hagemann-Jensen, Annika Krüger, Jilin Zhang, Christos Coucoravas, Ioannis Petrosian, Xian Xin, Ilhan Ceyhan, Joanna Rorbach, Dominic Wright, Björn Reinius

## Abstract

Sex-chromosome dosage represents a challenge for heterogametic species to maintain correct proportion of gene products across chromosomes in each sex. While therian mammals (XX/XY system) achieve near-perfect balance of X-chromosome mRNAs through X-upregulation and X-inactivation, birds (ZW/ZZ system) have been found to lack efficient compensation at RNA level, challenging the necessity of resolving major gene-dosage discrepancies in avian cells. Through allele-resolved multiome analyses, we comprehensively examined dosage compensation in female (ZW), male (ZZ), and rare intersex (ZZW) chicken. Remarkably, this revealed that females exhibit upregulation of their single Z through increased transcriptional burst frequency similar to mammalian X-upregulation, and that Z-protein levels are further balanced via enhanced translation efficiency in females. Global analyses of transcriptional kinetics elements in birds demonstrate remarkable conservation of the genomic encoding of burst kinetics between mammals and birds. Our study uncovers new mechanisms for achieving sex-chromosome dosage compensation and highlights the importance of gene-dosage balance across diverse species.

## Introduction

Vertebrate sex chromosome systems fall into two fundamental types, XX/XY and ZW/ZZ, defined by which sex is homogametic and which is heterogametic (*1*). In most mammals, including mice and humans, females are homogametic and possess two large, gene-rich sex chromosomes (XX), while males are heterogametic carrying one large and one degraded sex chromosome (XY). In ZW/ZZ systems, found in birds and reptiles, this relationship is inversed. While the systems evolved independently, they share being evolved from a once autosomal chromosome pair and the degeneration of the non-recombining sex chromosome defining the heterogametic sex by the process of “Muller’s ratchet” (*2, 3*), leaving the heterogametic sex (XY males and ZW females) with only a single copy of X/Z genes – thus unbalanced with the diploid autosomal gene expression network from which it originated. Pioneering theoretical work by Susumu Ohno (*4*) proposed that cells must restore such imbalance by a sex-chromosome-specific gene regulation mechanism. Today, it is experimentally well-characterized that mammals achieve dosage compensation at the transcriptional level by inactivating one X in females (XCI) and upregulating the single active X chromosome (XCU) in both sexes. Conversely, the question of Z chromosome dosage compensation remains debated to this day. Early studies in birds suggested little to no compensation (*5, 6*), while most recent works reported Male:Female Z-RNA-expression ratios of 1.2-1.6, suggesting inefficient, gene-specific, compensation rather than a chromosome-wide effect, thus questioning the generality of Ohno’s hypothesis (*6–10*). Some delimited segments of Z behave differently, such as the male-hypermethylated (MHM) regions (*11*) which display female-specific expression. More recently, a strongly male-biased Z-linked microRNA, miR-2954, has been reported to target dosage-sensitive Z-linked genes (*12–14*), which contribute to dosage compensation. However, current data on Z-chromosome dosage compensation remain largely fragmented, with partial biological information from different sources and no study to date providing allele-resolved single-cell data essential for understanding allelic expression dynamics central to this inquiry (*15*). Finally, while eukaryotic transcription is known to occur in stochastic bursts of RNA synthesis from the two alleles, transcriptional kinetics and its genomic encoding remain completely unexplored in birds.

Through a multimodal study, encompassing chromatin analyses, transcriptional kinetics, ribosomal profiling, and proteomics; using cell systems uniquely addressing avian dosage compensation at the allelic level, we uncover new insights and mechanisms of avian Z-chromosome dosage compensation.

## Results

To begin assessing Z-chromosome dosage compensation, we bred Red Junglefowl (RJF; *Gallus gallus*) and White Leghorn (WL; *G. g. domesticus*) chickens, as well as F1 offspring, and performed bulk RNA-sequencing on female and male tissues (brain, liver, kidney, skin, ovary, testis) (**Fig. 1A, fig. S1A**). We detected an average male-to-female Z-chromosome RNA level ratio of 1.57 across tissues (**Fig. 1B**) with sex-biased expression being strongly skewed to the sex chromosomes (**table S1**). Similar male bias was observed across the Z-chromosome (**Fig. 1C**), with exception of the ∼250kb-long MHM (*11*), known to harbour female-biased transcripts (*6, 16*). Z-linked genes retained from whole-genome duplication events in an ancient vertebrate ancestor (∼450 MYA) are thought to be more dosage sensitive (*17, 18*), and indeed, such genes showed lowered ratios in our RNA-seq data (∼1.36, P = 1.22 × 10^−7^, Tukey-HSD ANOVA corrected for tissue type) whereas more recent human orthologs (∼310 MYA)(*19*) or avian evolutionary strata (∼100 MYA)(*20*) did not differ across tissues (P_human_ = 0.96, P_avian_ > 0.11) (**fig. S1B**). Male-to-female rations were, overall, not associated with functional annotations in any of the tissues (FDR>0.05, GSEA Biological Processes).

**Figure 1.**
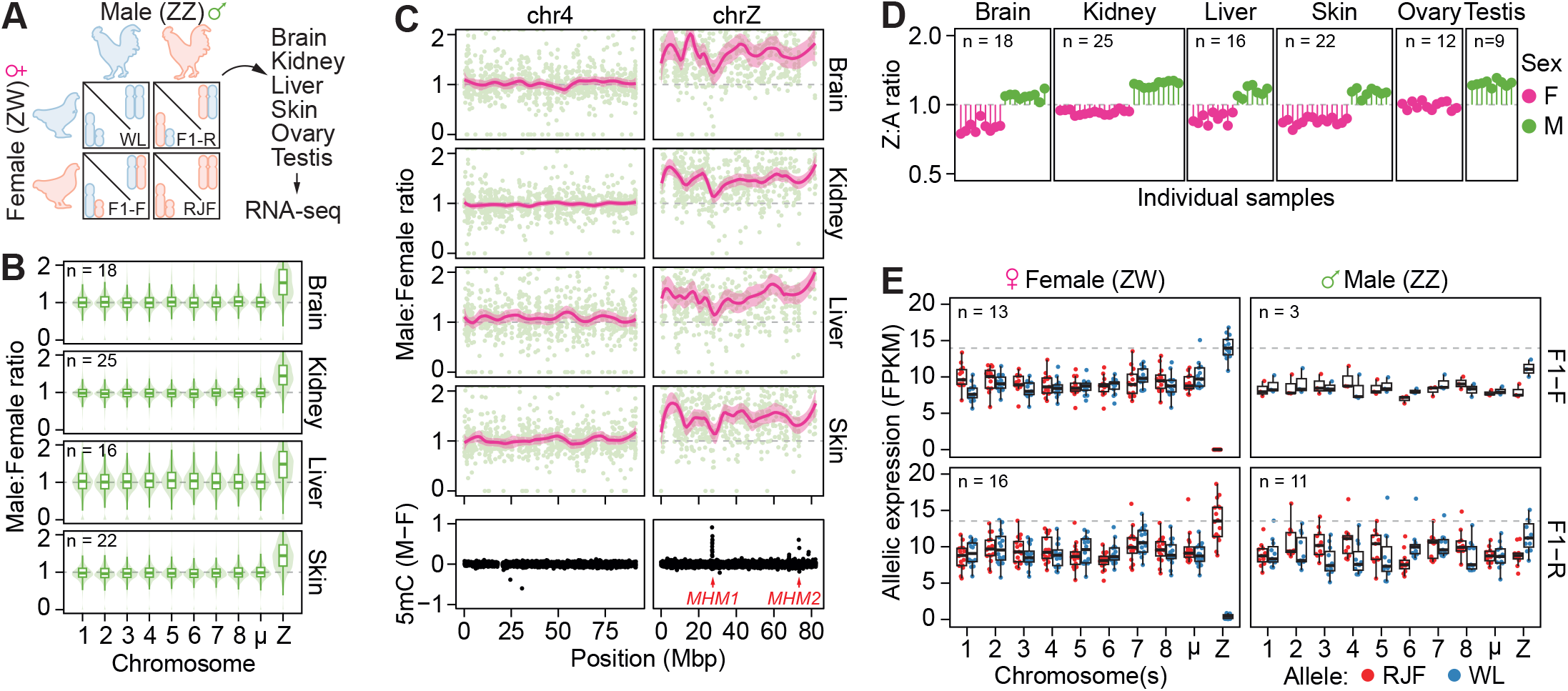
Z-chromosome upregulation of the female Z chromosome. **(A)** Schematic representation of the experimental set up. RNA from brain, kidney, liver, skin, ovary and testis tissues was isolated from purebred WL (White Leghorn; blue) or RJF (Red Junglefowl; red) or F1 crossbred chickens (F1-Forward: RJFmat x WLpat or F1-Reverse: WLmat x RJFpat) and used for allele-resolved RNA-sequencing using bulk UMI-Smartseq2. **(B)** Male:Female ratios of gene expression per tissue shown as boxplots over violin plots. Number of samples per tissue (n) shown in respective panels. Only expressed genes (average FPKM >1; chrA: n=384-1698, chrZ: n=542-607) were included in the analysis. µ denotes grouped micro-chromosomes 9-33. Data shown as median, first and third quartiles and 1.5x interquartile range (IQR). **(C)** Male:Female gene expression ratios along chromosomes 4 and Z per tissue. Average ratio per gene shown in green, with a rolling average (LOESS) ± 95% confidence interval shown in pink. On the bottom panel, Male:Female 5mC enrichment^49^ is shown for chromosomes 4 and Z, with male hypermethylated regions denoted in red. **(D)** Z:autosome ratios of bulk RNA-seq of WL, RJF and F1 chicken tissues. Each line and dot represent individual samples. Number samples per tissue shown in respective panels. Only expressed genes (FPKM >1; chrA: n=266-936, chrZ: n=351) were included in the analysis. Female and male samples coloured in pink and green respectively. **(E)** Boxplots of allelic expression (FPKM) in female and male samples for each chromosome for F1 tissue samples (see A.). RJF and WL alleles shown in red and blue respectively. µ denotes grouped micro-chromosomes 9-33. Number of samples (n) (average FPKM >1; chrA: n=266-936, chrZ: n=351) shown in respective panels. Data shown as median, first and third quartiles and 1.5x interquartile range (IQR).

Importantly, the observed male-to-female expression ratios (∼1.57) deviate from the 2-fold ratio expected in complete absence of Z-dosage compensation between ZZ (male) and ZW (female) genotypes. In line with this, comparing Z-linked expression to diploid autosomes (AA) per sample showed that female Z expression (Z:AA) was higher than expected for all tissues (**Fig. 1D**). To explore Z-dosage compensation at allele-specific level, we called genetic variants from the pure WL and RJF parental breeds allowing allelic expression analyses in F1 offspring. After filtering (see **“materials and methods”**), we retained 193,130 allele-informative variants covering 83.64% of expressed genes across all chromosomes, enabling high-resolution allelic inference (**fig. S1C**). Whereas there was a slight bias towards the WL Z chromosome in F1 males compared to autosomes (**fig. S1D**), we did not observe an overrepresentation of differentially expressed Z-linked genes in pure WL breeds compared to RJF (OR = 1.05, P = 0.724, Fisher’s exact test, **table S2**). Interestingly, utilizing our allelic expression measurements, we observed that the single Z chromosome in all female tissues was distinctly upregulated both compared to autosomes and to the separate transcriptional output of each male Z allele **(Fig. 1E, fig. S1E)**, indicating that partial dosage compensation is achieved through hyperactivation of the female Z chromosome. Intriguingly, this observed Z-chromosome upregulation resembles mammalian XCU earlier found in mammals (*21–24*).

To extensively characterise Z-upregulation, we derived primary F1 chicken embryonic fibroblast (CEF) cell lines from eggs of RJF/WL intercrosses and performed a comprehensive array of omics profiling (**Fig. 2A**). During analysis, we noticed that one CEF line expressed chromosome W in addition to two Z alleles, which we resolved as triploid ZZW intersex by DNA-seq and karyotyping (**fig. S2A-E**). This is a naturally viable genotype, albeit exceedingly rare (0.1-0.5%), resulting from chromosomal nondisjunction during oogenesis (*25*) **(fig. S2F)**. This unexpected individual allowed us to compare the degree of Z-linked dosage compensation between diploid males (ZZ:AA), diploid females (ZW:AA), and triploid intersex (ZZW:AAA) genotypes, thereby uncoupling potential effects of W on dosage compensation. Expression relative to autosomes revealed that Z-linked expression was partially compensated in both ZW and ZZW genotypes (**fig. S3A**), suggesting an unexpected flexibility in avian dosage compensation. Interestingly, whereas RNA-seq of CEF lines reconfirmed Z-upregulation in ZW females on par with *in vivo* tissues (mean fold-change 1.54, **Fig. 2B**), dosage compensation in the ZZW intersex individual was not mediated through Z-upregulation but buffering of autosomal expression (**fig. S3B**), as has previously been observed for other species (*26*). On the transcriptional level, intersex CEFs show similar gene expression patterns as females, e.g. with high expression of W-linked *HINTW* and *SPINW*, believed to be involved in ovarian development, and female-biased Z-linked *BHMT2* expression **(table S3)**, in agreement with observations that ZZW intersex chicks are phenotypically similar to females until week 20 post hatching (*27*).

**Figure 2.**
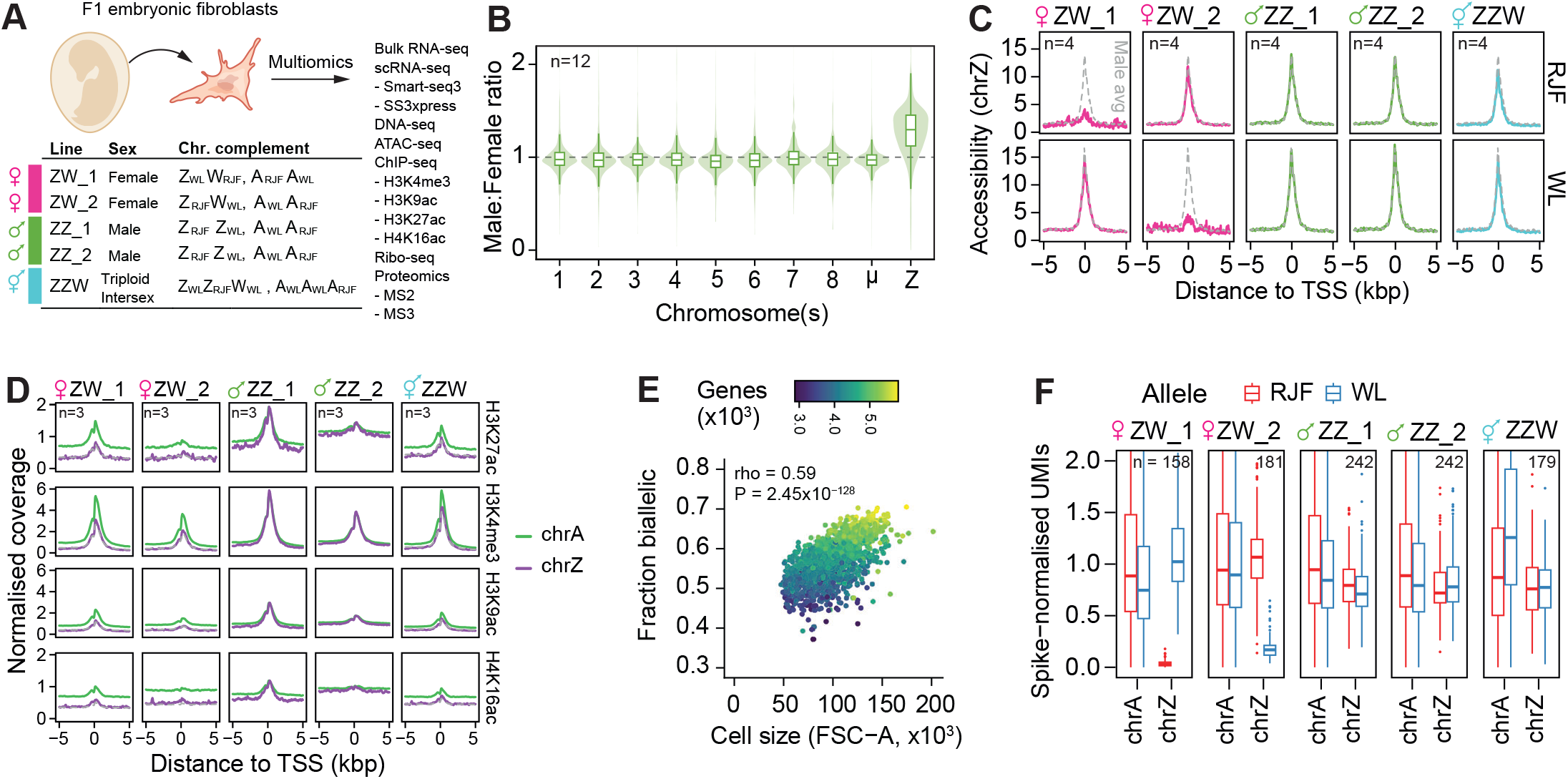
Chromatin states and transcriptional kinetics of Z-chr upregulation. **(A)** Schematic representation of the experimental set up. Primary chicken embryonic fibroblasts (CEF) were isolated from 10-13-day old F1 chicken embryos and used for multi-omic characterization. **(B)** Boxplots of Male:Female gene expression ratios per chromosome for chicken embryonic fibroblasts (CEFs). µ denotes grouped micro-chromosomes 9-33. Number of samples (n) shown in respective panels. Only expressed genes (average FPKM >1; chrA: n=352-1325, chrZ: n=351) were included in the analysis. Data shown as median, first and third quartiles and 1.5x IQR. n=12 defined as: CEF lines derived from n=2 female and n=2 male embryos grown as 3 independent replicates (n=4*3=12). **(C)** Density plots of allele-resolved ATAC-seq signal enrichment at transcription start sites (TSS), per sample and allele, represented as means ± 95% CI obtained from 4 independent replicates. Female, male and intersex samples shown in pink, green and teal respectively. **(D)** Density plots of quantitative ChIP-seq signal enrichment around transcription start sites (TSS), per histone modification and sample, represented as means ± 95% CI obtained from 3 independent replicates. Autosomal signal shown in green and Z-chr signal in purple. **(E)** Scatterplot of number of biallelically expressed genes (average FPKM >20, n=1033-3579 genes) per cell (n=1382), by cell size (FSC-A) based on FACS and single-cell RNA-seq (Smart-seq3) data. Point colours are denoted as genes with FPKM>1 per cell. **(F)** Boxplots of allele-resolved spike-in normalised UMI counts of single-cell RNA-seq (Xpress-seq) of chicken embryonic fibroblasts (CEFs) per sample and allele. Number of single cells (n) shown in the respective panels. Only expressed genes (average FPKM >1, chrA: n=8467, chrZ: n=424) were included in the analysis. Data shown as median, first and third quartiles and 1.5x IQR.

To gain further insights into the regulation of Z-upregulation, we mapped chromatin accessibility (ATAC-seq) on the CEF lines. Similar to our previous findings of XCU in mice (*24*), chromatin accessibility was not increased on the upregulated Z allele (**Fig. 2C** and **fig. S4A-C**), suggesting that Z-upregulation is primarily controlled at the transcriptional level. Using transcription factor footprinting analysis, we investigated the differential binding of transcription factors between sexes, via pairwise comparisons of Z and autosomes for all three sexes. FOX and GATA transcription factor families showed preferential binding on male and female chromosomes respectively, both on Z and autosomes (**fig. S4D-E, table S4**). Interestingly, E-box (enhancer box) transcription factors were enriched on female and intersex Z chromosomes, and not on the male Z, a Z-specific difference not detected in the autosomes (**fig. S4D-E, table S4**). We next performed multiplexed quantitative ChIP-seq (*28*) (EpiFinder) for four permissive histone modifications (H3K4me3, H3K9ac, H3K27ac and H4K16ac) associated with promoter- and enhancer features (**fig. S5A-B**). Although these modifications were overall associated with gene expression levels (**fig. S5C**), they were not enriched with Z-upregulation (**Fig. 2D**).

The intersex line and allelic resolution allowed us to explore the MHM region from a new angle. As expected, we detected MHM-expression in ZW females and the lack thereof in ZZ males, and intriguingly, intersex ZZW displayed accessible chromatin and biallelic RNA expression (**fig. S6**), implying that the MHM region is controlled by the presence of the W rather than Z-chromosome copy numbers, as previously suggested (*11*).

To explore Z-upregulation at cellular allelic regulation, we performed Smart-seq3 (*29*) scRNA-seq full-transcript-coverage deep-sequencing on the CEF lines (**fig. S7A**), reconfirming Z-chromosome upregulation within individual female cells (**fig. S7B**) and biallelic expression of the MHM region in intersex ZZW cells (**fig. S6C**). Single-cell resolution also allowed us to investigate how general cell-intrinsic features associate with RNA expression output in birds, where cell size was highly correlated with genes detected (rho = 0.6, P = 3.47×10^−133^) and number of RNA molecules per cell (rho = 0.5, P = 3.03×10^−66^) (**fig. S7C**). Furthermore, cell size was also associated with the fraction of genes expressed biallelically due to stochastic allelic transcription (rho = 0.59, P = 1.05×10^−132^) (**Fig. 2E**), indicating that the same principal laws of cell scaling and random monoallelic expression apply in birds as in mouse and human (*30, 31*).

To enable direct comparison between diploid and triploid expression levels we performed Xpress-seq, a method for full-length scRNA-seq developed from Smart-seq3xpress (*32*) including UMI-containing exogenous spike-in RNA (*33*), allowing precise counting of original mRNA molecules in single cells for all five CEF lines (**fig. S8**). Notably, this re-confirmed female-specific Z-upregulation and that ZZW lacked Z-upregulation (**Fig. 2E**). At the single-cell level, eukaryotic transcription is inherently stochastic and occurs in short bursts of activity from individual alleles (*34, 35*). Transcriptional kinetics can be encoded as burst frequency (rate of transcription pulses) and size (average number of molecules produced during a burst) using a two-state telegraphic model of transcription (*36*). We previously showed that X-upregulation in mouse is driven by increased transcriptional burst frequency, but not increased burst size (*23, 24*), and here sought to characterize transcriptional kinetics in birds for the first time. To this end, we inferred parameters of transcriptional bursting for each allele in the Xpress-seq data (see **“materials and methods”** and **fig. S9A-B**). First, we established the principles of burst kinetics in chicken which mirrored what is observed in humans and mice (*36, 37*) (**Fig. 3A**). Specifically, we found burst size to be driven by the presence of TATA-box in the core promoter, with initiator elements having a small additive effect (**Fig. 3B**), consistent with the notion that TATA promoters show high rates of continuous transcription (*38, 39*). Conversely, burst frequency was associated with cis-regulatory activity through permissive histone modifications (**fig. S9C-D**), as has previously been shown in mice (*40*), suggesting a universal control of burst kinetics in mammals and birds, and that the genomic encoding of bursting is deeply conserved in vertebrate species. We next explored the kinetic modulus of Z-upregulation, observing that burst frequency was increased on Z relative to autosomes in female ZW cells (P_WL-allele_ = 2.68 × 10^−4^, P_RJF-allele_ = 0.029, MWU Test) whereas Z alleles did not in male ZZ and intersex ZZW cells (RJF alleles analysed in triploid cells) (**Fig. 3A**). Conversely, burst size remained close to autosomal levels for the same comparisons (P>0.41) (**fig. S10A**). It should be noted that inference of transcriptional kinetics is only robust for individual alleles (*36*), necessitating allelic scRNA-seq data herein provided, and allowing accurate kinetic inference of RJF alleles but not WL in the triploid line.

**Figure 3.**
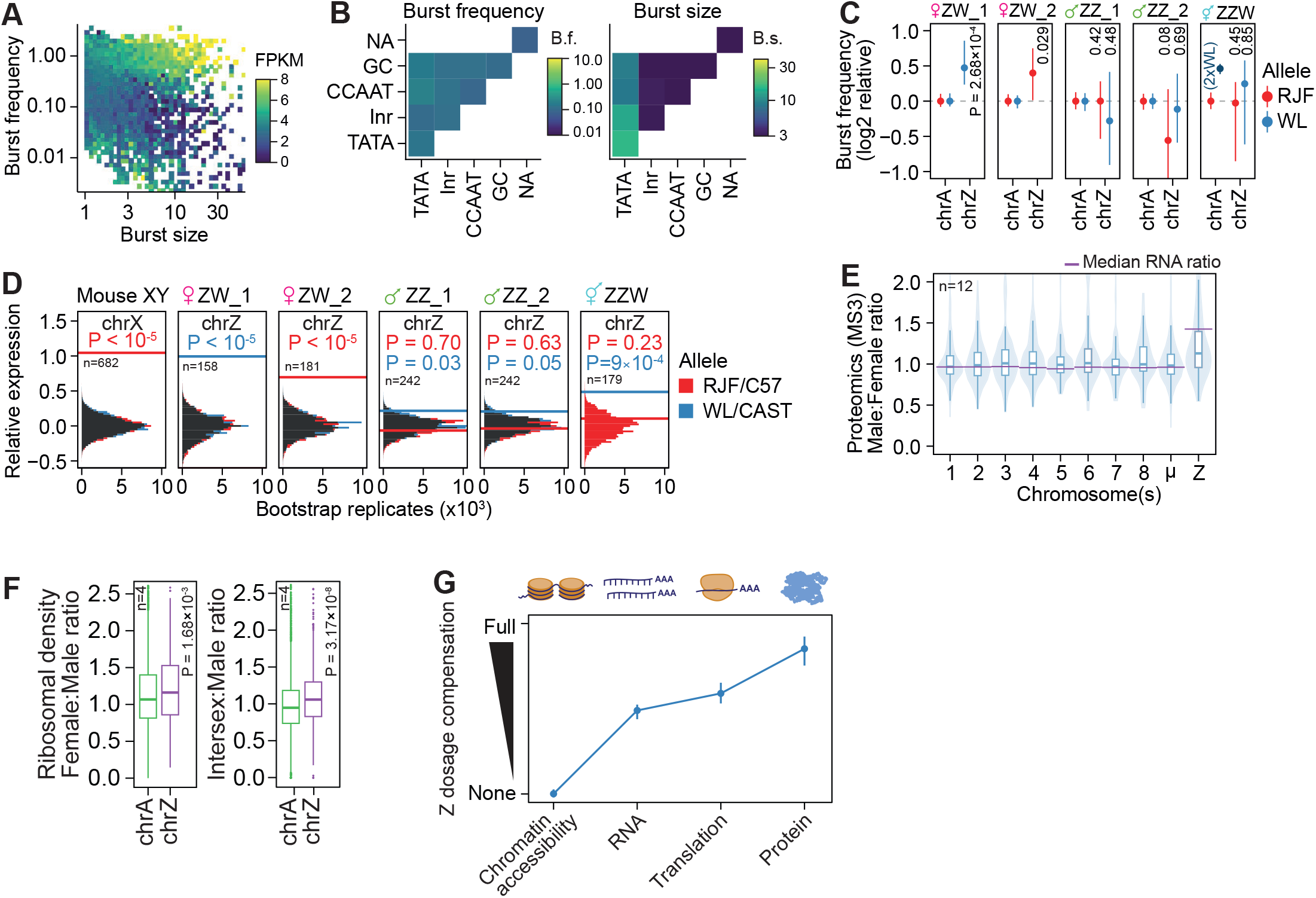
Transcriptional and post-transcriptional control of Z-upregulation. **(A)** Scatter plot of burst frequency (y-axis) and burst size (x-axis) inferred for expressed genes (7392 genes, average FPKM > 1) in CEFs, coloured based on FPKM-normalised expression level. **(B)** Heatmaps of core promoter elements and their correlation with burst frequency (left) or burst size (right) for expressed genes (7392 genes, average FPKM >1). **(C)** Allele-resolved, log2-relative burst frequency per cell for autosomal (n=4586) and Z-linked (n =229) genes, plotted as median ± 95% CI. RJF and WL alleles shown in red and blue, respectively. P-values from MWU tests between respective chrA alleles or against RJF for ZZW. **(D)** Relative expression levels of autosomal and X-linked genes in mouse and Z-linked genes in chicken. Histogram of medians of relative expression in FPKM of randomly subsampled genes, compared to median of chrX for mouse (chrA genes: n=10025, chrX genes: n=276) or chrZ for chicken (chrA genes: n=7180, chrZ genes: n=332), based on scRNA-seq in mouse fibroblasts50 and CEFs, with number of cells used shown in black. P-values denote empirical bootstrap P-values at n=105. C57 (C57BL6/J) and CAST (CAST/EiJ) denotes mouse alleles. **(E)** Boxplot over violin plots of Male:Female ratios of protein abundances (MS3) per chromosome (chrA: n=62-895; chrZ: n=90) in CEFs. Purple vertical lines indicate median Male:Female ratios of gene expression based on bulk RNA-seq in CEFs. Data shown as median, first and third quartiles and 1.5x interquartile range (IQR). µ denotes grouped micro-chromosomes 9-33. Number of samples (n) defined as: CEF lines derived from n=2 female and n=2 male embryos grown as 3 independent replicates (n=4*3=12). **(F)** Female:Male and Intersex:Male ratios of translation efficiency, calculated as gene expression-normalised ribosome-protected footprint (RPF) FPKM counts for autosomes and chrZ. Only expressed genes were included in the analysis (RPF > 1 & RNA FPKM > 1; chrA genes: n=9213-9220; chrZ genes: n=443). MWU tests were used for significance testing. Data shown as median, first and third quartiles and 1.5x interquartile range (IQR). **(G)** Schematic representation of a multi-layered model of Z-chromosome dosage compensation in birds. At the genomic level, the single female Z chromosome does not display enhanced accessibility. At the gene expression level, compensatory transcriptional upregulation of the female Z is apparent, and driven by increased transcription burst frequency. Z-linked transcripts display higher translational efficiency, partially contributing to an overall rebalancing and near-complete dosage compensation between ZZ males and ZW females at the proteomic level.

Given the similarities in transcriptional response of Z-upregulation in chicken to X-upregulation in mice, we sought to compare the two directly. To compensate for gene content differences between chromosomes and species, we utilised a bootstrapping approach (*24*) to compare gene-level transcriptional metrics. The degree of upregulation was highly similar in chicken (ZW) and mouse (XY) fibroblasts (**Fig. 3D, fig. S10B**), suggesting a similar mechanism of transcriptional upregulation between evolutionary unrelated sex chromosomes.

Despite the apparent tolerance to large Z-linked RNA expression differences between sexes, sex chromosome aneuploidies are embryonic lethal in chicken (*41, 42*). To understand how these seemingly contradictory concepts resolve at the proteomic level, we performed tandem mass spectrometry for the CEF lines using both MS2 and MS3 spectra (see **“materials and methods”**). Through an additional round of protein fractionation (MS3), we were able to improve detection sensitivity and identified up to 2,122 proteins per line including up to 91 Z-encoded proteins (**fig. S11A**). Protein abundances showed good agreement with RNA-seq expression (Spearman’s Rho = 0.55-0.57, P < 1.7×10^−112^) (**fig. S11B**) Interestingly, while Male:Female ratios were significantly increased for Z also at the proteome levels (1.2 fold, P = 7.63×10^−7^, MWU test) the Z-level difference were diminished compared to RNA level (P = 1.45×10^−5^, one-sample Mann Whitney U-test) (**Fig. 3E** and **fig. S11C-F**), indicating a second layer of dosage compensation in birds. Indeed, gene-wise Male:Female ratios shifted towards RNA expression for Z-linked genes (**fig. S11F**), suggesting that ZW samples produce more Z-encoded protein per mRNA expression unit.

Seeking to explore whether the effect observed on the protein level is due to differences in translational efficiency, we performed ribosome profiling (Ribo-seq) on the five CEF lines. Using ribosome-protected fragment counts normalised to RNA expression for each gene (see **“materials and methods”** and **fig. S12A-C**), we calculated translation efficiency (TE) rates for autosomes and the Z-chromosome in the three sexes. Indeed, female and intersex samples displayed higher TE rates for the Z chromosome than for autosomes compared to males (P_ZW/ZZW_ < 7.53 × 10^−4^, P_ZZ_ > 0.1, MWU) (**fig. S12D**), suggesting increased ribosome occupancy on female and intersex Z-linked transcripts. To compare ribosome occupancies in a gene-wise manner, we calculated Female:Male and Intersex:Male ratios for expressed genes and across different expression cutoffs, which further confirmed significantly higher TE for the female and intersex Z chromosomes (P_F:M_ < 0.0017, P_I:M_ < 3.17 × 10^−8^, MWU) (**Fig. 3F**). Although the detected effect was modest, it is consistent with reports of an increased translational efficiency of X-linked transcripts in eutherian mammals (*43, 44*), again highlighting shared mechanisms of Z/X-upregulation. Notably, while female-to-male RNA levels are near-perfectly balanced in mouse and human, X:AA ratios are not (*23, 24*), explaining the need of translational upregulation in mammals carrying one active X chromosome per cell similar to ZW females.

Our findings do not exclude that additional mechanisms are at play in avian dosage compensation. miR-2954 is known to target a subset of dosage-sensitive Z-linked mRNAs in males (*12–14*), thus we explored to what degree miR-2954 might explain dosage compensation in our data. miR-2954 quantification by RT-qPCR showed 4-14-fold higher expression in male CEFs and tissues compared to females, as expected, while revealing intermediate levels in the ZZW genotype (**fig. S13A-B, table S5**). Stratifying genes containing or lacking miR-2954 target sequence (29% CEF-expressed Z genes containing target; Methods), we indeed found lowered Male:Female ratios for miR-2954 targets on Z but not autosomes, while intersex comparisons suggest that ZZW behave more like males than females in this aspect (**fig. S13C-D, table S5**). Thus, while miR-2954 provide partial relief, we conclude that chicken rely on a multi-layered dosage compensation strategy also involving control of transcriptional burst frequency and transitional rates.

## Discussion

In this study, we aimed to uncover the mode and degree of dosage compensation in chicken. Using bulk transcriptomics on pure-line and allele-resolved chicken tissues, we demonstrate that sex chromosome dosage compensation is achieved through chromosome-wide transcriptional upregulation of the single Z allele in females. We investigated transcriptional burst kinetics using high-sensitivity, allele-resolved single-cell transcriptomics on primary chicken fibroblasts, establishing that Z-upregulation is driven by increased burst frequency, but not size, resembling the kinetic mode of the evolutionarily distinct X-upregulation observed in mouse (*23, 24*). Despite the ∼310 MYA separating birds and mammals, we additionally found that the core promoter elements controlling transcriptional burst size remain the same, with TATA and Inr promoter elements correlating with increased burst size, suggesting that these molecular mechanisms may be fundamental across vertebrates.

Chromatin accessibility and permissive histone modifications have been previously suggested to at least partially underlie X-chromosome upregulation in mouse (*45–47*). However, despite a pronounced compensation on the transcriptional level, using allele-resolved ATAC-seq and quantitative ChIP-seq, we observed no differences in accessibility or chromatin state between female and male Z-chromosomes, in line with previous reports of non-linear relationships between chromatin states and gene expression in X-upregulation (*24, 45, 46*).

The inclusion of a natural but rare triploid intersex chicken (ZZW) enabled us to uncouple the effect genetic sex may have on Z-upregulation. Despite similarities between female and intersex samples, we found no evidence of Z-chromosome upregulation in the ZZW genotype. Conversely, we observed that incomplete dosage compensation in intersex cells was mediated through the dampening of autosomal expression. Thus, Z-upregulation by increased transcriptional bursting in female cells results from carrying only one Z allele, not the presence of W.

Despite transcriptional dosage compensation, protein stoichiometries are ultimately crucial for cellular and organismal fitness and survival. We found that the transcriptional upregulation of the female Z chromosome is further boosted at the proteome level, achieving a significant dosage rebalancing between males and females. Our proteomic data are in line with that of a recent report (*48*). Our Ribo-seq data indicates that this is least partially driven by higher ribosomal density, and thus increased translational efficiency of Z-transcripts in females. Intriguingly, similarly Z-enriched ribosomal density was observed in ZZW intersex cells suggesting the involvement of W-linked factors. Increased translation efficiency of Z in female, but not male, cells contrasts to the situation in mammals, where X translation is elevated in both sexes (*43, 44*). However, while X-linked RNA levels are nearly balanced between female and males the transcriptional upregulation of the their single active X (XaXi / XaY) does not match the biallelic RNA levels of autosomes (*23, 24*). Consequently, enhanced Z-translation rate on top of transcriptional Z-upregulation in ZW females is analogous to the scenario in mammalian cell across sexes.

Together, our study uncovered a complex interplay of transcriptional and translational mechanisms that synergistically achieve extensive dosage compensation of the avian Z chromosome, and an unexpected similarity to mammalian dosage compensation when all regulatory layers are coherently taken in consideration.

## Acknowledgements

We thank all members of the Reinius lab for their input, and employees of Epigenica for early access and technical support related to EpiFinder. Protein identification and quantification were carried out at the Proteomics Biomedicum core facility at Karolinska Institutet and Xpress-seq library preparation was performed at Xpress Genomics, Stockholm, Sweden. Sequencing was performed at Xpress Genomics and National Genomics Infrastructure (NGI), SciLifeLab Stockholm, Sweden.

## Author contributions

NP performed the CEF culturing and analyses, including FACS analyses, bulk and single-cell RNA-seq, karyotyping, RT-qPCRs, proteomics, analysed data, prepared figures, and wrote the manuscript. AL performed data pre-processing, QC and allelic analysis of ATAC-seq, bulk tissue RNA-seq and CEF scRNA-seq, analysis of burst kinetics, and FACS data, prepared figures, and wrote the manuscript. SW performed ATAC-seq and ChIP-seq analyses. MHJ generated Smart-seq3 libraries. AK and JR performed Ribo-seq. JZ and XX performed analysis of tissue RNA-seq data. IP performed footprinting analysis. IC performed karyotyping. CC generated tissue RNA-seq libraries. DW conceived the study, planned and conducted chicken crossing, and performed tissue dissection. BR conceived and supervised the study, secured funding, derived the CEF lines, analysed data, and wrote the manuscript.

## Data and code availability

Raw and processed data is available through ArrayExpress under accessions E-MTAB-14391 (DNA-seq), E-MTAB-14390 (ATAC-seq), E-MTAB-14392 (ChIP-seq), E-MTAB-14393 (Ribo-seq). ArrayExpress accession numbers pending for bulk RNA-seq, scRNA-seq, and proteomics, and will be available upon peer-reviewed publication of this work. Code to reproduce this work is available at github (https://github.com/reiniuslab/Z-upregulation).

## Supplementary Materials

Figs. S1-S13

Materials and Methods

Tables S1-S5

## Supplementary Materials

**Figure S1.**
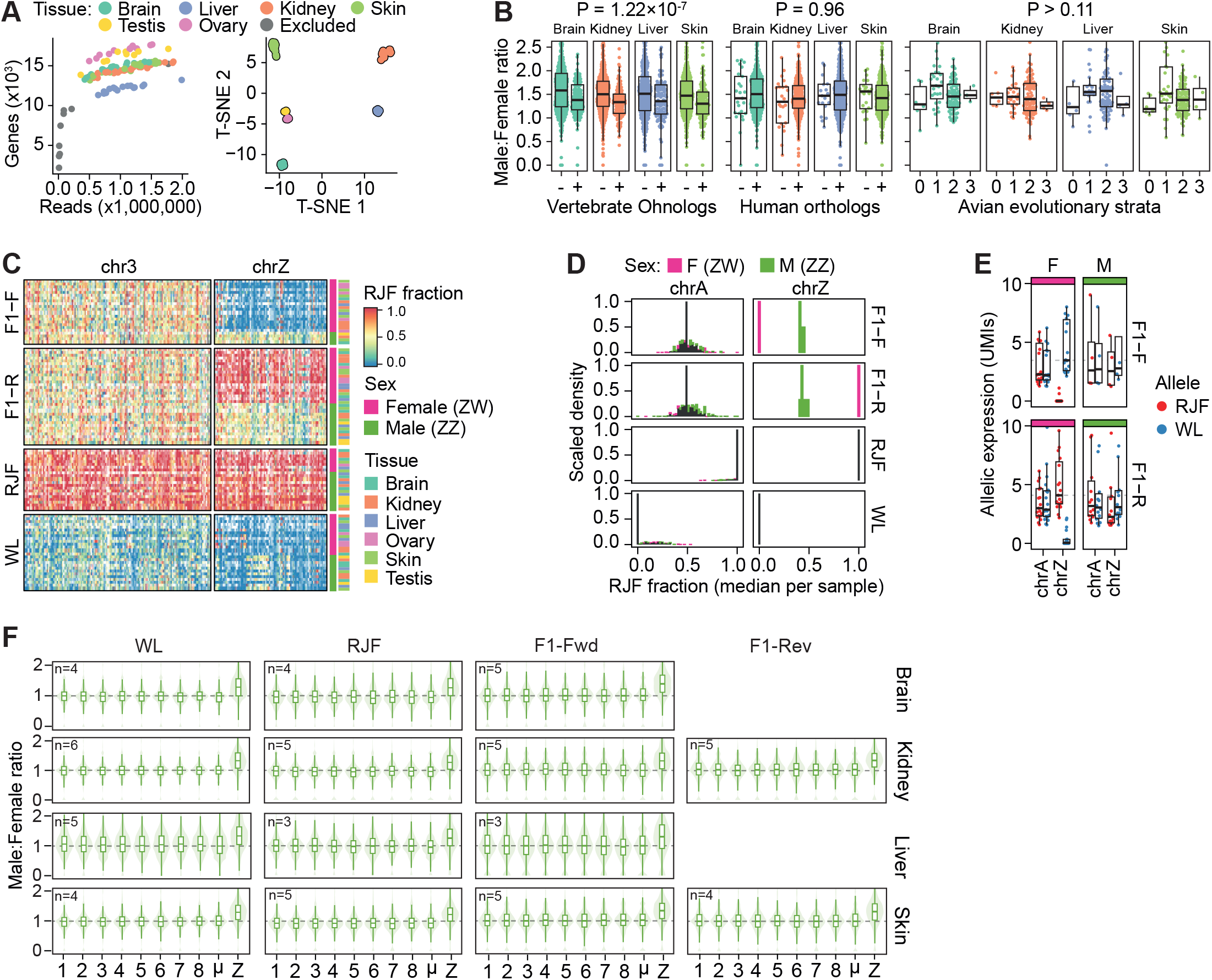
Allele-resolved bulk RNA-seq reveals transcriptional upregulation of the female Z chromosome. **(A)** Quality assessment of bulk RNA-seq libraries of tissue samples. On the left: Scatterplot of the number of thousands of genes detected (y-axis) and the number of sequencing reads in millions (x-axis), with each point representing an independent RNA-seq library. Different colours represent libraries prepared from distinct tissue samples (Brain: n=18, Kidney: n = 25, Liver: n=16, Skin: n=22, Ovary: n=12, Testis: n=9) with samples excluded due to low quality represented in grey. On the right: t-distributed stochastic neighbour embedding (t-SNE) of RNA-seq libraries, with each tissue cluster represented in different colours. **(B)** From left to right: Boxplots of Male:Female gene expression ratios of bulk RNA-seq of chicken tissues for vertebrate ohnologs (671 genes) (+) and non-ohnologs (-), human orthologs (450 genes) (+) and non-orthologs (-) and Z-linked genes belonging to different evolutionary strata (0 = oldest, 3 = newest; 133 genes). Tukey-HSD ANOVA was used for significance testing. **(C)** Heatmap of allelic expression for chromosome 3 (389 genes) and chromosome Z (234 genes) for pure (WL = White Leghorn; n=26, RJF = Red Junglefowl; n=21) and forward (F1-F, RJF x WL; n=22) and reverse (F1-R, WL x RJF; n=33) F1-derived tissue samples (total: n = 102). **(D)** Histogram of median fraction of allelic red Junglefowl (RJF) reads of bulk RNA-seq of pure (WL; White Leghorn or RJF; Red Junglefowl) or F1 (F: RJF x WL, R: WL x RJF) female and male tissue samples shown as scaled density. **(E)** Allelic expression in UMI counts (derived from UMI-containing reads) of bulk UMI-containing RNA-seq (SS2-UMI) for F1 reciprocal cross male and female tissue samples (F1-Forward; RJF x WL; female: n=18, male: n=4; F1-Reverse; WL x RJF; female: n=19, male: n = 14). RJF (Red Junglefowl) and WL (White Leghorn) alleles shown in red and blue respectively. Data shown as median, first and third quartiles and 1.5x IQR. **(F)** Male:Female ratios of gene expression per tissue and cross shown as boxplots over violin plots. Number of samples per tissue (n) shown in respective panels. Only expressed genes (average FPKM >1) were included in the analysis (chr1: n=1683-1785, chr2: n=1127-1178, chr3: n = 1018-1065, chr4: n=931-977, chr5: n = 807-832, chr6: n = 434-480, chr7: n = 421-450, chr8: n = 424-452, chrµ: n = 5195-5594, chrZ: n=596-624). µ denotes grouped micro-chromosomes 9-33. Data shown as median, first and third quartiles and 1.5x interquartile range (IQR).

**Figure S2.**
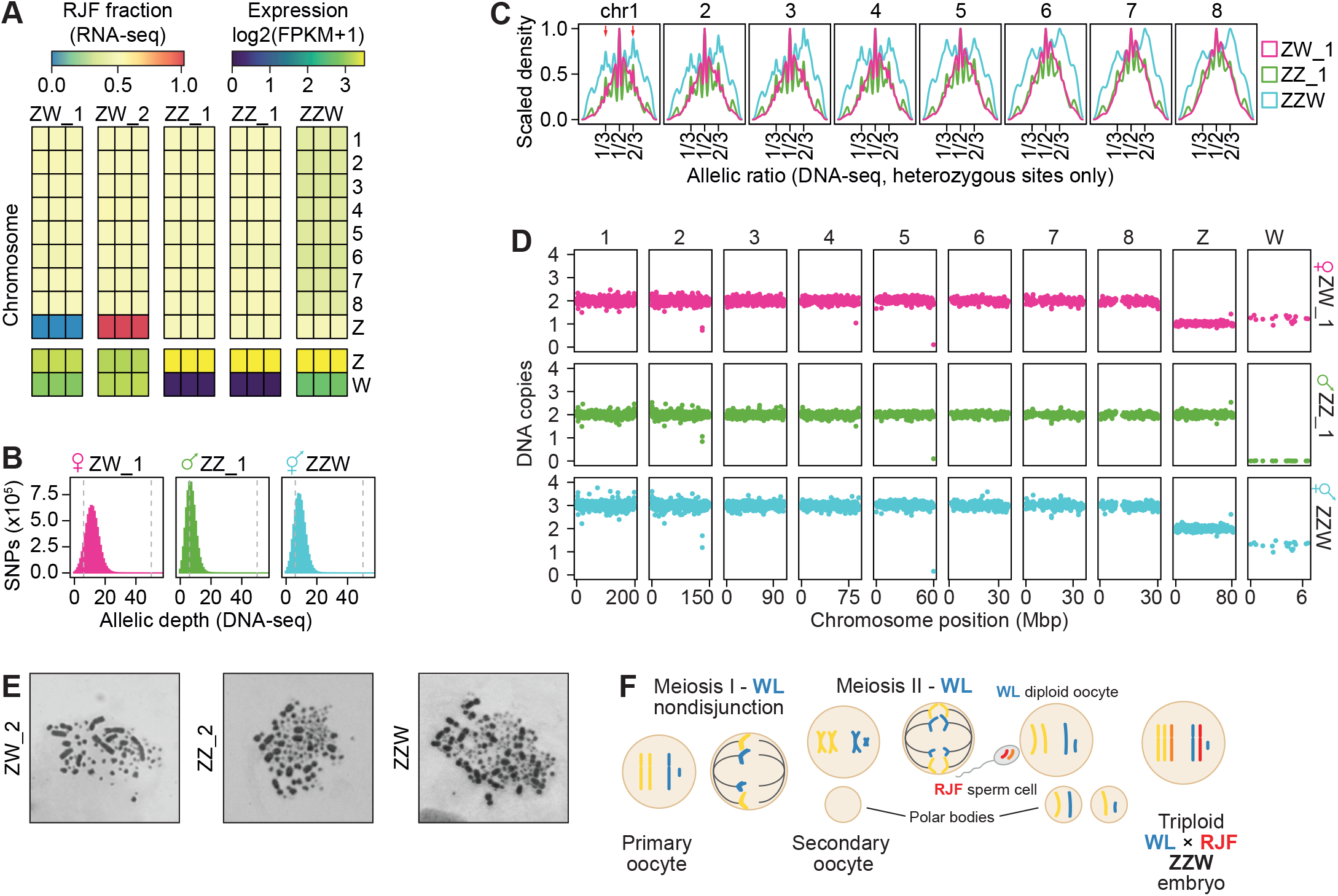
Identification of a triploid intersex (ZZW) sample. **(A)** Allele expression heatmap for bulk RNA-seq for chromosomes 1-8, Z and W (total of 2943 genes: chr1=629, 2=439, 3=396, 4=376, 5=313, 6=195, 7=173, 8=182, Z=240) shown per sample (N=15 from 5 primary CEF lines * 3 independent replicates), coloured by fraction of red Junglefowl (RJF) reads (top) and log2-normalised reads per kilobase million-normalised reads counts for the Z and W chromosomes (bottom). **(B)** Histogram plot of number of single-nucleotide polymorphisms (SNPs) detected on the y-axis and allelic coverage of DNA-sequencing libraries on the x-axis. Vertical lines denote cutoffs used. **(C)** Scaled density of allelic ratios over heterozygous sites based on DNA-sequencing data for female, male and intersex samples shown in pink, green and teal respectively, for chromosomes 1-8. For diploid F1 samples, allelic ratios are expected at 0.5. Red arrows indicate the unequal distribution of allelic ratios due to the presence of two WL allele copies and one RJF allele copy per chromosome in ZZW intersex samples. **(D)** Scatter plots of DNA copy number (y-axis) per chromosomal position in Mbp (x-axis) shown for (macro)chromosomes 1-8, Z and W. Female, male and intersex samples shown in pink, green and teal respectively. **(E)** Representative metaphase spreads depicting the karyotypes of a female (ZW_2; n=15 identifiable macrochromosomes), male (ZZ_2; n=16 identifiable macrochromosomes) and intersex (ZZW; n=24 identifiable macrochromosomes) samples. **(F)** Schematic representation of the formation of a triploid intersex embryo. During meiosis I, a nondisjunction event occurring at anaphase I in the primary oocyte can lead to the aggregation and transfer of sister chromatids to a single secondary oocyte, generating a void polar body. Meiosis II proceeds to generate two haploid polar bodies and one diploid oocyte which upon fertilisation by a haploid sperm cell generates a triploid zygote.

**Figure S3.**
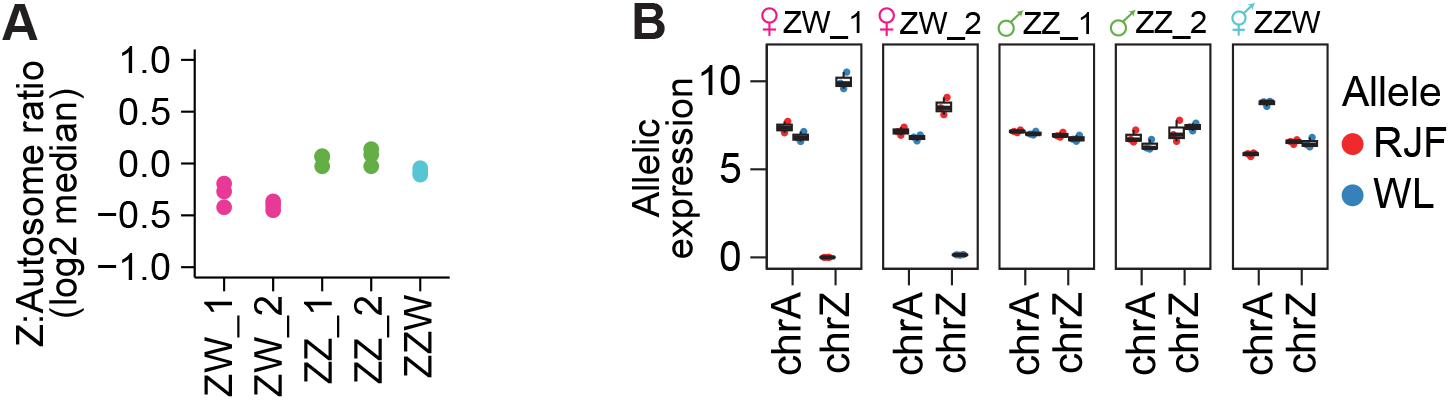
Amplification-free bulk RNA-seq confirms Z-chromosome upregulation of the female Z. **(A)** Log2-normalised median Z:autosome ratios of gene expression (FPKM>1, total number of genes used in analysis: 10361; autosomal genes = 9888, Z-linked genes=473) per sample (N=15 from 5 primary CEF lines * 3 independent replicates) of bulk RNA-seq (Truseq) in CEF lines. Female, male and intersex samples shown in pink, green and teal respectively. **(B)** Boxplots of allelic gene expression in FPKM of bulk RNA-seq (Truseq). N=15 from 5 primary CEF lines * 3 independent replicates. Allele-resolved genes included in the analysis = 4956 (average FPKM>1, autosomal genes = 4737, Z-linked genes = 219). RJF and WL alleles shown in red and blue respectively.

**Figure S4.**
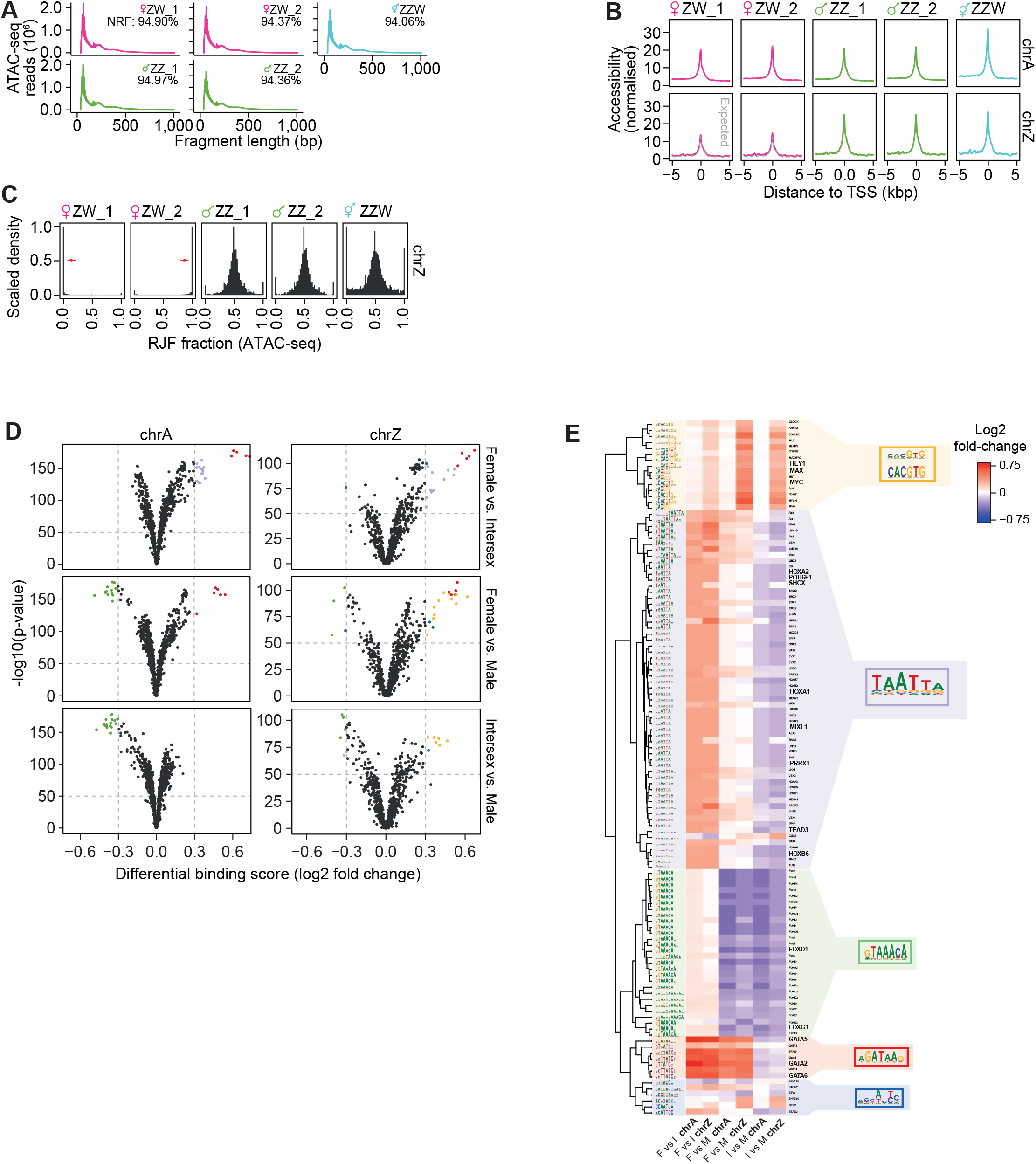
Chromatin accessibility and transcription factor footprinting of the chicken Z chromosome. **(A)** ATAC-seq library fragment size distribution with fragment length in base pairs on the x-axis and library read numbers on the y-axis, shown as mean of n=4 independent replicates. Female, male and intersex samples shown in pink, green and teal respectively. NRF = non-redundant fraction. **(B)** Density plots of normalised accessibility for autosomes (top panel) and Z-chromosome (bottom panel) per primary CEF line (mean of n=4 replicates) with distance from the TSS in kilo base pairs on the x-axis. Female, male and intersex samples shown in pink, green and teal respectively. **(C)** Density plots of allelic chromatin accessibility shown as fraction of RJF (red Junglefowl) allelic reads for chromosome Z per primary CEF line. Mean of n=4 independent replicates. Female, male and intersex samples shown in pink, green and teal respectively. Red arrows shown to indicate allelic skew of chrZ in ZW samples. **(D)** Volcano plots of the differentially bound transcription factors between sexes for autosomes (chrA) and for chrZ. The x-axis shows the log2 fold change of the differential binding score as calculated by the TOBIAS BINDetect module, and the y-axis represents the significance of the observed change. The most significant transcription factors (>|0.3| change in binding score) for each comparison have been coloured, whereas FOX and GATA transcription factor families, which consistently showed preferential binding on male and female chromosomes respectively, were marked with different colour. Interestingly, a group of transcription factors was detected preferentially bound on both female chrZ and intersex chrZ when compared to the male chrZ, a difference that was not detected in the autosomes. Note that for motifs recognized by more than ten different possible transcription factors, their names are not shown. **(E)** Heatmap showing the log2 fold change of the differential binding scores of the transcription factors per comparison (Left to right: Female - Intersex: autosomes, Female - Intersex: ChrZ, Female - Male: Autosomes, Female-Male: ChrZ, Intersex-Male: Autosomes and Intersex-Male: ChrZ). Only transcription factors with differential binding score >|0.3| in at least one comparison were included. The colour indicates the differential binding score, with positive score (red) suggesting preferential binding to the first sex of the comparison (female/intersex) and negative score (blue) indicating preferential binding to the second sex of the comparison (male/intersex). The transcription factors were clustered based on their motif similarity and consensus motifs were created per cluster. The colour-coding corresponds to transcription factors belonging to the same family or recognizing the same motif (shown as seqlogos).

**Figure S5.**
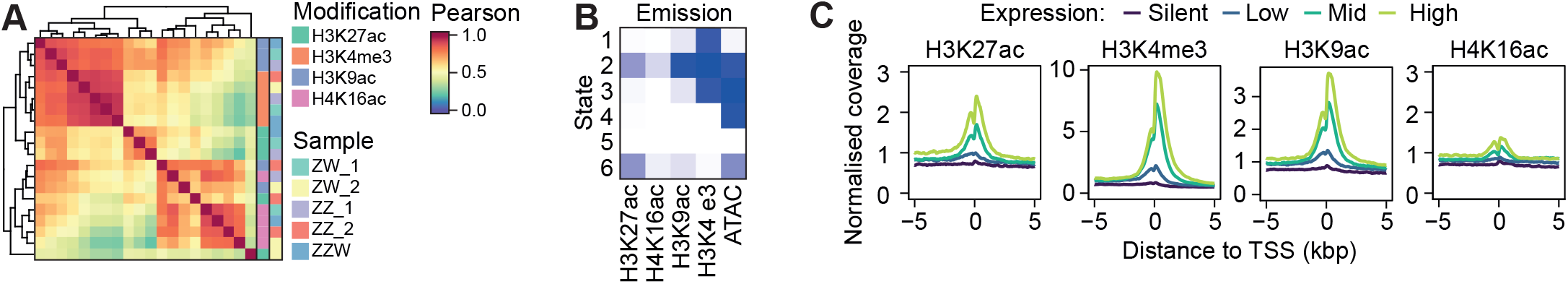
Quantitative ChIP-seq of the chicken Z chromosome. **(A)** Pearson correlation heatmap of quantitative ChIP-seq histone modifications and samples with hierarchical clustering shown on the right for n=20 samples (5 CEF lines * 4 histone modifications) based on n=71,262 10kb bins. **(B)** Heatmap of chromHMM’s emission parameters with each row corresponding to a different chromatin state and each column to a different chromatin mark (quantitative ChIP-seq) or chromatin accessibility (ATAC-seq). The colour gradient shows the probability of observing the respective chromatin mark or accessibility in that state, with darker colours corresponding to higher probability and lighter colours to lower probability. **(C)** Density plots of normalised coverage of quantitative ChIP of CEFs around genomic transcription start sites (TSS) in kilo base pairs per histone modification, based on n = 3 independent replicates. The different colours correspond to levels of gene expression (silent, low, mid, high), based on bulk RNA-seq in CEFs.

**Figure S6.**
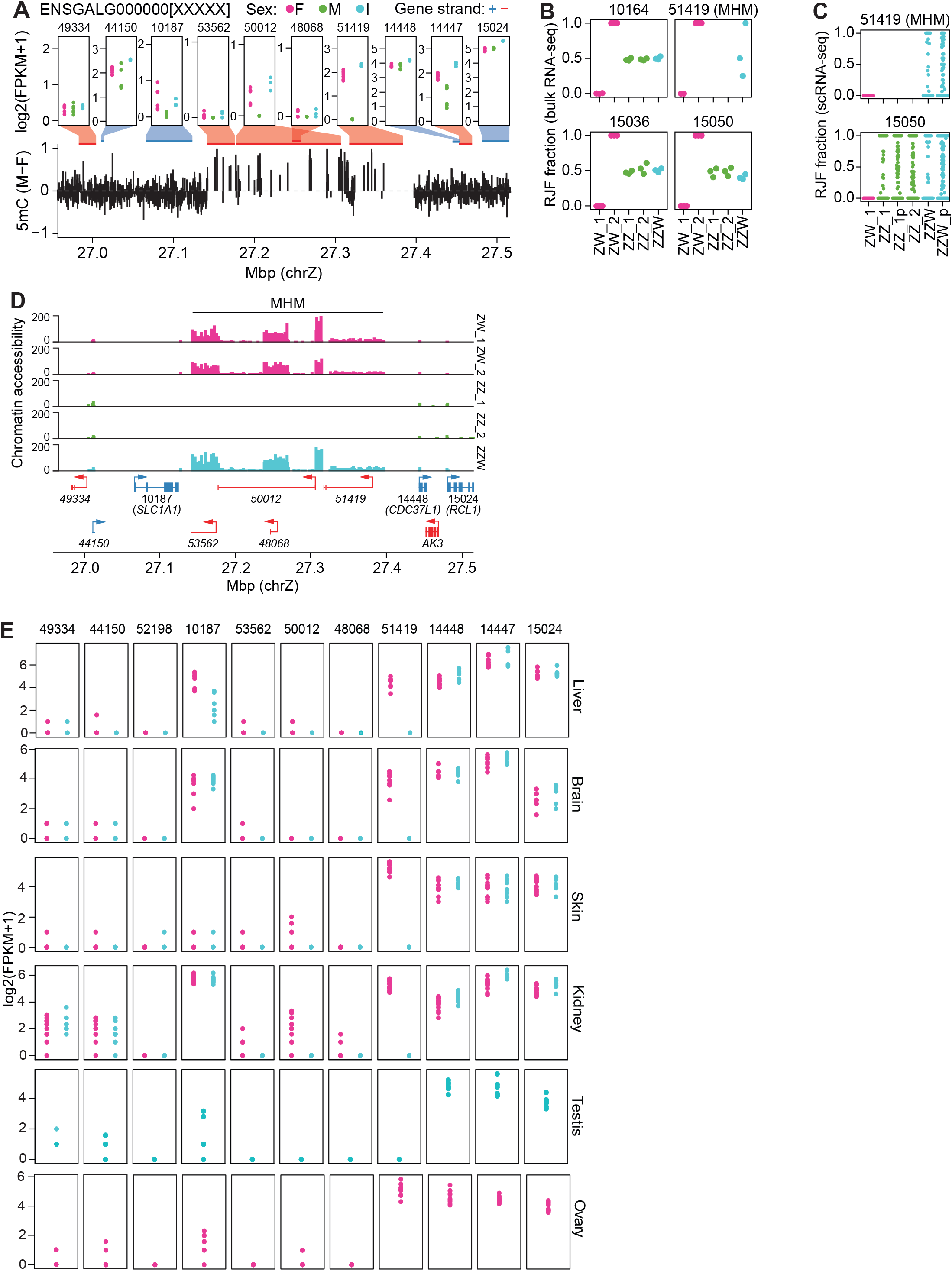
The male hypermethylated (MHM) region is accessible and expressed in females and intersex. **(A)** Top: Log2-normalised gene expression (in fragments per kilobase million, FPKM) of transcripts of the Z-linked male hypermethylated region (MHM) in female, male and intersex samples (n=3 independent replicates) based on bulk RNA-seq in CEFs. Transcript names shown as “ENSGALG000000[XXXXX]” with the last five digits corresponding to Ensembl transcript IDs appearing above each transcript’s panel. Bottom: Male:Female 5mC enrichment49 in the MHM region. **(B)** Allele-resolved expression of transcripts of the MHM region of chrZ displayed as fraction of Red Junglefowl (RJF) reads for each sample based on bulk RNA-seq. Female, male and intersex samples shown in pink, green and teal respectively. **(C)** Allele-resolved single-cell RNA-seq shown as fraction of Red Junglefowl (RJF) reads for transcripts of the MHM region for each sample based on scRNA-seq (Smart-seq3) of CEFs (number of cells: ZW_1: n=329, ZZ_1: n=323, ZZ_1p: n=350, ZZ_2: n=366, ZZW: n=364, ZZW_p: n=350, where “_p” denotes CEFs of earlier passage). **(D)** Genomic tracks of bulk chromatin accessibility for the Z-linked male hypermethylated (MHM) region [27.1-27.4 Mbp] grouped by sample and coloured by sex, with female, male and intersex samples shown in pink, green, and teal respectively. **(E)** Log2-normalised gene expression (in fragments per kilobase million, FPKM) of transcripts of the Z-linked male hypermethylated region (MHM) in female and male samples based on bulk RNA-seq in WL, RJF and F1 tissues. Transcript names shown as “ENSGALG000000[XXXXX]” with the last five digits corresponding to Ensembl transcript IDs appearing above each transcript’s panel.

**Figure S7.**
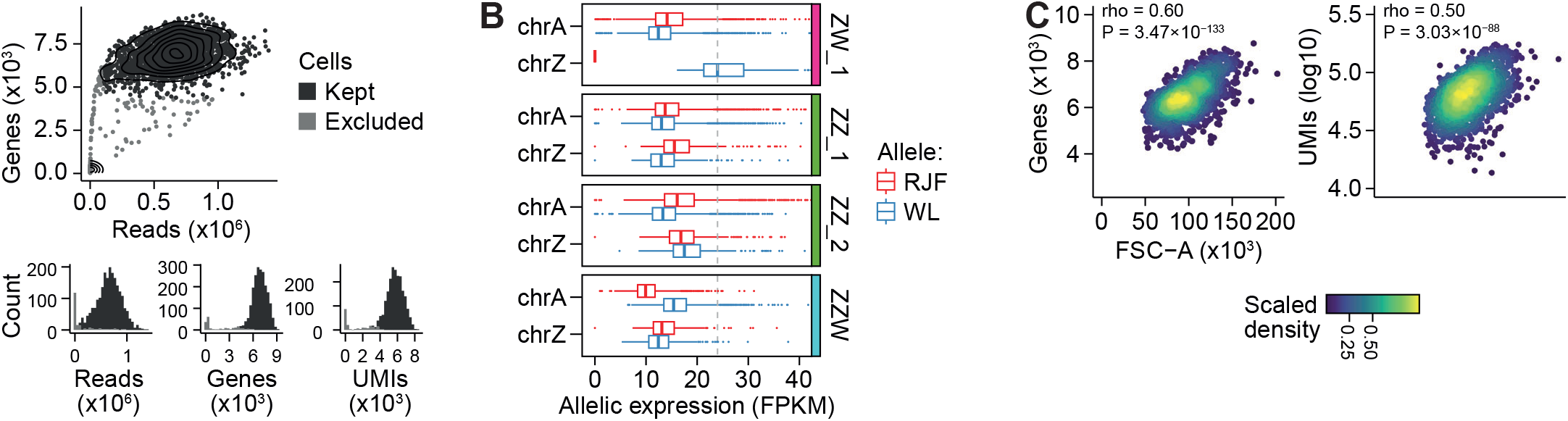
Transcriptional burst kinetics of the Z chromosome using Smart-seq3. **(A)** Quality assessment of single-cell RNA-seq libraries (Smart-seq3). Top: Scatterplot of number of genes detected (in thousands; n=4397-9197 kept genes) on the y-axis in relation to number of sequencing reads (in millions) per sequenced cell. Cells excluded due to low quality shown in grey (kept: n = 2082, excluded: n=200). Bottom: Number of read counts by number of sequencing reads in million (left), detected genes in thousands (middle) and unique-molecular identifiers (UMIs) in thousands (right). **(B)** Boxplots of allelic gene expression in FPKM for autosomes (chrA; n=5649 genes) and the Z chromosome (chrZ; 283 genes) based on single-cell RNA-seq (Smart-seq3) for each CEF line (n=329-364 cells). Grey dashed line denotes allelic expression levels of the single female Z chromosome in sample ZW_1. Data shown as median, first and third quartiles and 1.5x IQR. **(C)** Left: Scatterplots of number of genes expressed (in thousands) per cell, by cell size (FSC-A) based on FACS and single-cell RNA-seq (Smart-seq3) data. Right: Scatterplot of number of UMIs detected per cell (in log10) in relation to cell size (FSC-A). Colour gradient denotes scaled density. Statistics represents Spearman correlation tests.

**Figure S8.**
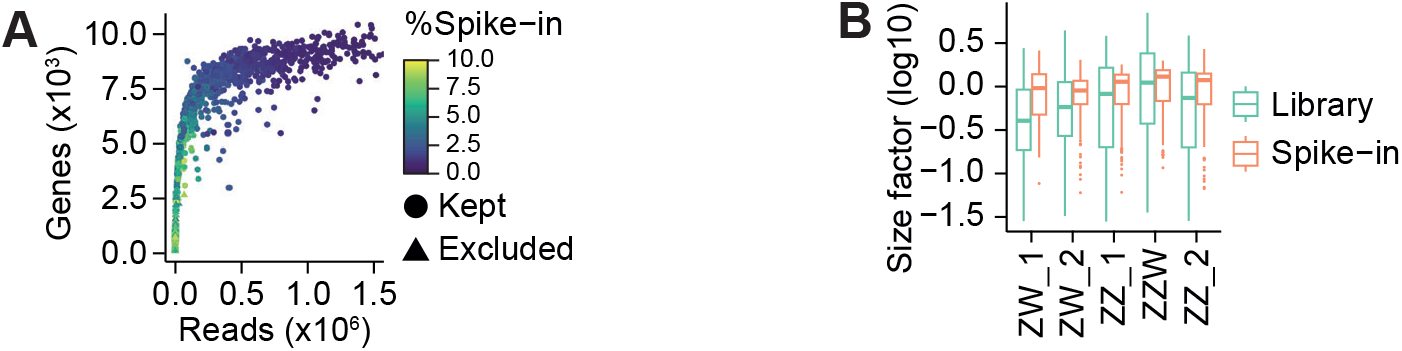
Spike-in normalised single-cell RNA-sequencing. **(A)** Scatterplot of number of genes detected in thousands (y-axis) relative to number of sequencing reads per cell in million (x-axis) based on spike-in single-cell RNA-seq (Xpress-seq). The percentage of spike-ins per cell is denoted by the colour gradient. Cells not passing quality filtering shown as triangles (n kept = 1002, n excluded = 195). **(B)** Boxplot of size factor normalisation (in log10) of spike-in single-cell RNA-seq libraries for each CEF line. Data shown as median, first and third quartiles and 1.5x IQR.

**Figure S9.**
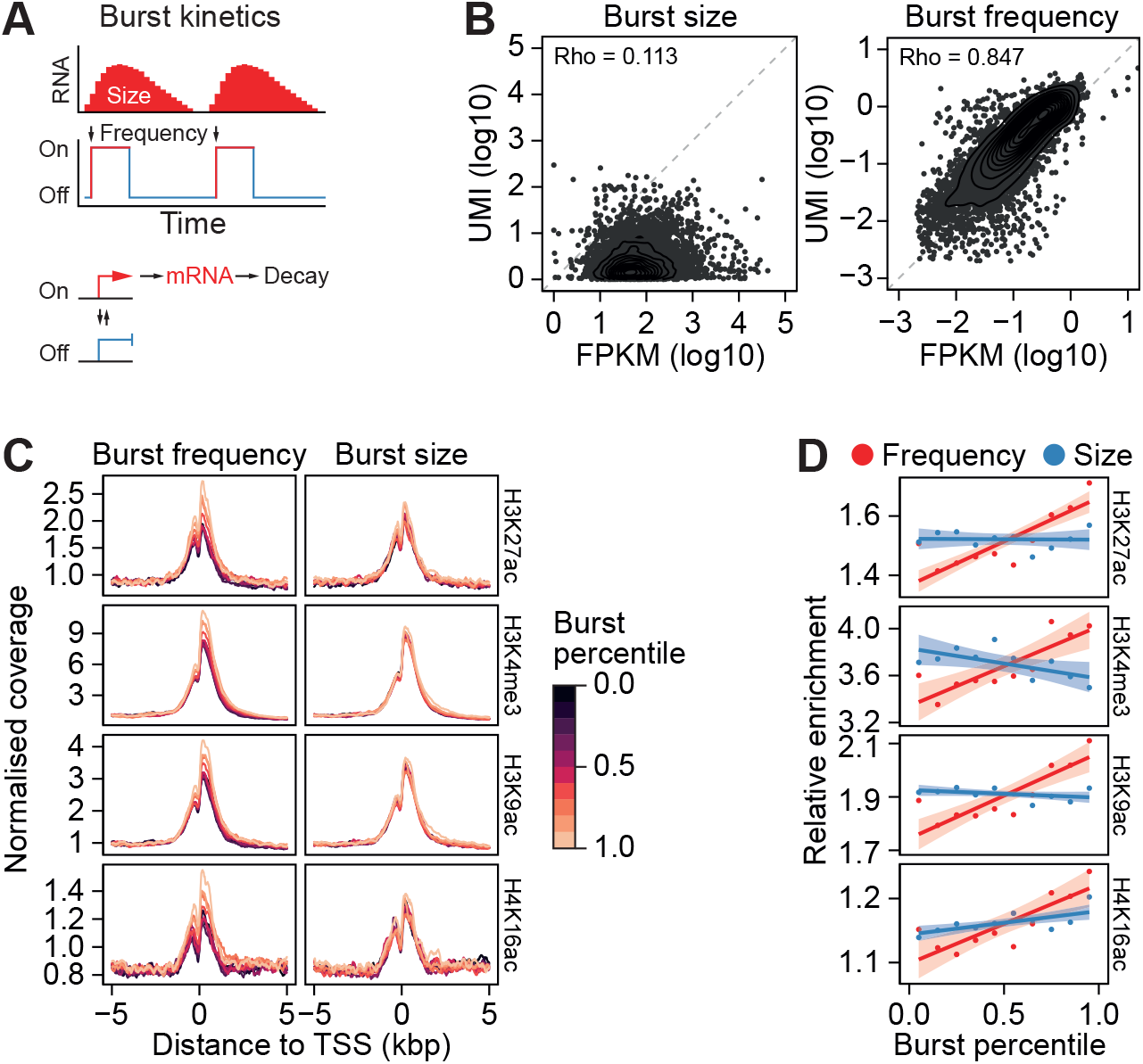
Transcription burst frequency is associated with permissive histone modifications. **(A)** Schematic representation of transcriptional burst kinetics. During each bursting event, RNA molecules are produced. Top: Transcription burst size represented as number of RNA molecules produced during each transcription burst event (also termed “on” state). Diagram of burst frequency with state on the y-axis (“on” denoting active transcription, “off” denoting no transcription). Bottom: Simplified schematic depicting the transcription burst parameters used to model transcription burst kinetics. **(B)** Correlation of UMI-containing reads and FPKM-normalised counts for burst size and burst frequency based on spike-in normalised single-cell RNA-seq (Xpress-seq). Left: Scatter plot showing the correlation between unique molecular identifier (UMI) counts (log10) and fragment per kilobase million (FPKM)-normalised single-cell RNA-seq counts in log10 scale for burst size. Right: Scatter plot showing the correlation between unique molecular identifier (UMI) counts and fragment per kilobase million (FPKM)-normalised single-cell RNA-seq counts in log10 scale for burst frequency. Spearman correlation coefficients (rho) displayed in respective panels. **(C)** Quantitative ChIP enrichment around genomic TSS, with distance from TSS in kilo base pairs on the x-axis for each histone modification in CEFs, relative to transcription burst dynamics. Colour gradient denotes the burst percentile, with lighter colours denoting higher degree of burst frequency (left) or size (right), obtained from spike-in single-cell RNA-seq (Xpress-seq) data in CEFs. **(D)** Line plots of correlation between quantitative ChIP relative enrichment of each histone modification and transcriptional burst frequency (in red) or burst size (in blue), expressed as burst percentile, obtained from spike-in normalised single-cell RNA-seq (Xpress-seq) in CEFs. Data shown as linear model mean and ± 95% CI.

**Figure S10.**
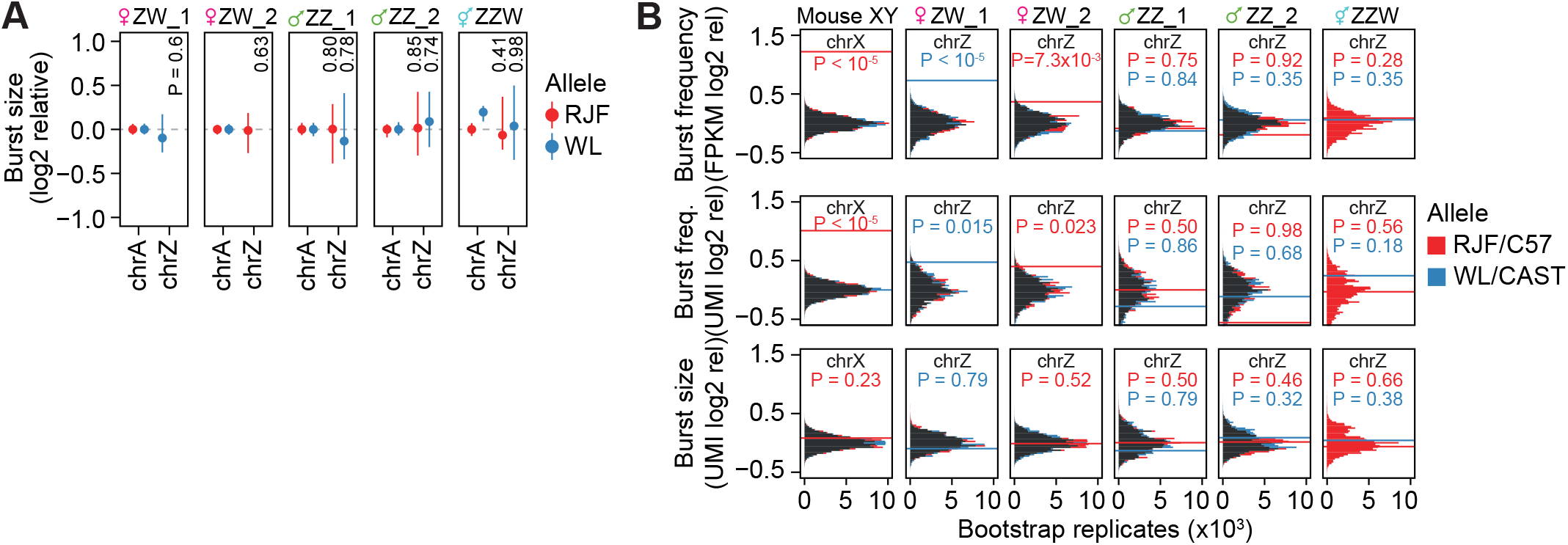
Transcriptional burst kinetics of the chicken Z chromosome mechanistically resemble mammalian X chromosome upregulation. **(A)** Log2-relative allele-resolved transcription burst size of autosomes (chrA) and the Z-chromosome (chrZ) per CEF line, obtained from spike-in normalised single-cell RNA-seq (Xpress-seq). P-values are empirical P-values from bootstrap resampling with n=105. Data shown as median ± 95% C.I. **(B)** Comparison of transcription burst frequency and size between autosomes and chrX in mouse (chrA genes: n=10025, chrX genes: n=276) or chrZ for chicken (chrA genes: n=7180, chrZ genes: n=332). Histogram of medians of relative burst frequency in FPKM (top) or UMI counts (middle) and burst size in UMI counts (bottom) of randomly subsampled genes, compared to median of chrX for mouse or chrZ for chicken, obtained from single-cell RNA-seq in mouse fibroblasts (n = 682 cells) and CEFs (ZW_1: n=158, ZW_2: n=181, ZZ_1: n=242, ZZ_2: n=242, ZZW: n=179). Number of permutations (bootstrap replicates) shown on x-axis. Red colour denotes C57BL6/J allele in mouse comparisons and RJF in chicken comparisons and blue colour denotes CAST/Eij allele for mouse comparisons and WL for chicken comparisons.

**Figure S11.**
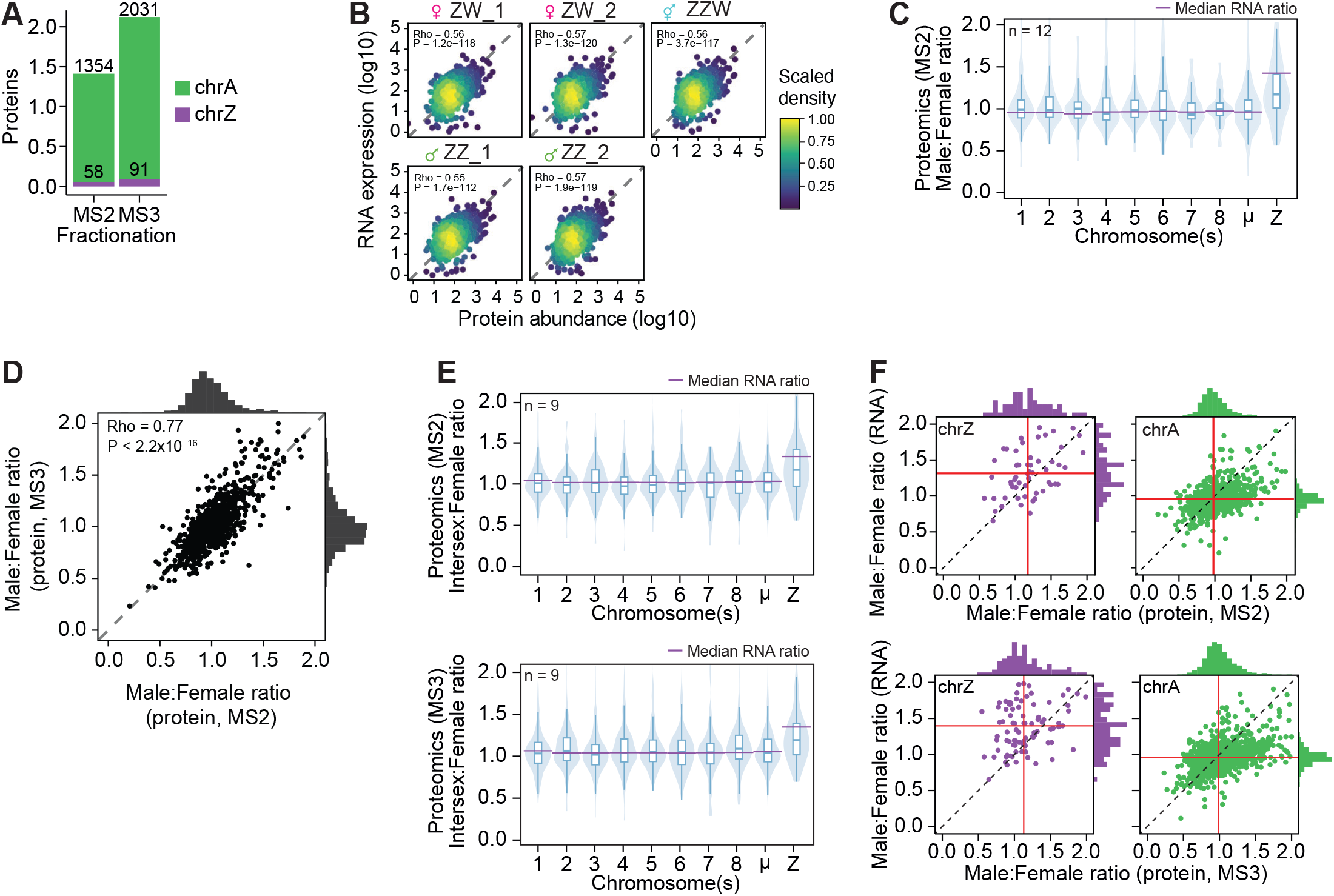
Tandem mass spectrometry reveals significant dosage compensation of the Z chromosome. **(A)** Bar plot displaying the number of unique autosomal (in green) and Z-linked (in purple) proteins (y-axis) identified in two rounds of fractionation using tandem mass spectrometry (MS2) or triple-stage mass spectrometry (MS3) in CEFs. **(B)** Scatter-plots of RNA expression (y-axis) and protein abundance (x-axis) obtained from amplification-free bulk RNA-seq (Truseq) and tandem mass spectrometry (MS2) data in CEFs, per sample, coloured by scaled density. Spearman correlation shown at the top. **(C)** Violin and boxplots of Male:Female ratios of protein abundances (MS2: number of proteins per chr: chr1:n =174; chr2: n=109; chr3: n=100; chr4: n=97; chr5: n=100; chr6: n=49; chr7: n=55; chr8: n=49; chrµ: n=621; chrZ: n=58) per chromosome in CEFs, displayed as mean of n=3 independent replicates. Purple vertical lines indicate median Male:Female ratios of gene expression based on amplification-free bulk RNA-seq in CEFs. Data shown as median, first and third quartiles and 1.5x interquartile range (IQR). **(D)** Scatterplot of Male:Female ratios of protein abundances based on MS2 (x-axis) and MS3 (y-axis) measurements for proteins detected in both MS2 and MS3 datasets (n = 1002), with histograms of distributions shown for MS3 (y-axis) and MS2 (x-axis). Spearman correlation shown over the scatter. **(E)** Violin and boxplots of Intersex:Female ratios of protein abundances based on MS2 (top; number of proteins per chr: chr1:n =174; chr2: n=109; chr3: n=100; chr4: n=97; chr5: n=100; chr6: n=49; chr7: n=55; chr8: n=49; chr9-33: n=621; chrZ: n=58) and MS3 (bottom; number of proteins per chr: chr1:n =264; chr2: n=162; chr3: n=158; chr4: n=134; chr5: n=146; chr6: n=74; chr7: n=79; chr8: n=68; chr9-33: n=945; chrZ: n=91) per chromosome in CEFs, displayed as mean of n=3 independent replicates. Purple vertical lines indicate median Intersex:Female ratios of gene expression based on bulk RNA-seq (Truseq) in CEFs. Data shown as median, first and third quartiles and 1.5x interquartile range (IQR). **(F)** Scatterplots of Male:Female ratios of gene expression based on amplification-free bulk RNA-seq in CEFs (Truseq) on y-axis and protein abundance measurements using MS2 (top; chr1:n =174; chr2: n=109; chr3: n=100; chr4: n=97; chr5: n=100; chr6: n=49; chr7: n=55; chr8: n=49; chrµ: n=621; chrZ: n=58) or MS3 (bottom; number of proteins per chr: chr1:n =264; chr2: n=162; chr3: n=158; chr4: n=134; chr5: n=146; chr6: n=74; chr7: n=79; chr8: n=68; chr9-33: n=945; chrZ: n=91) for detected proteins and expressed genes.

**Figure S12.**
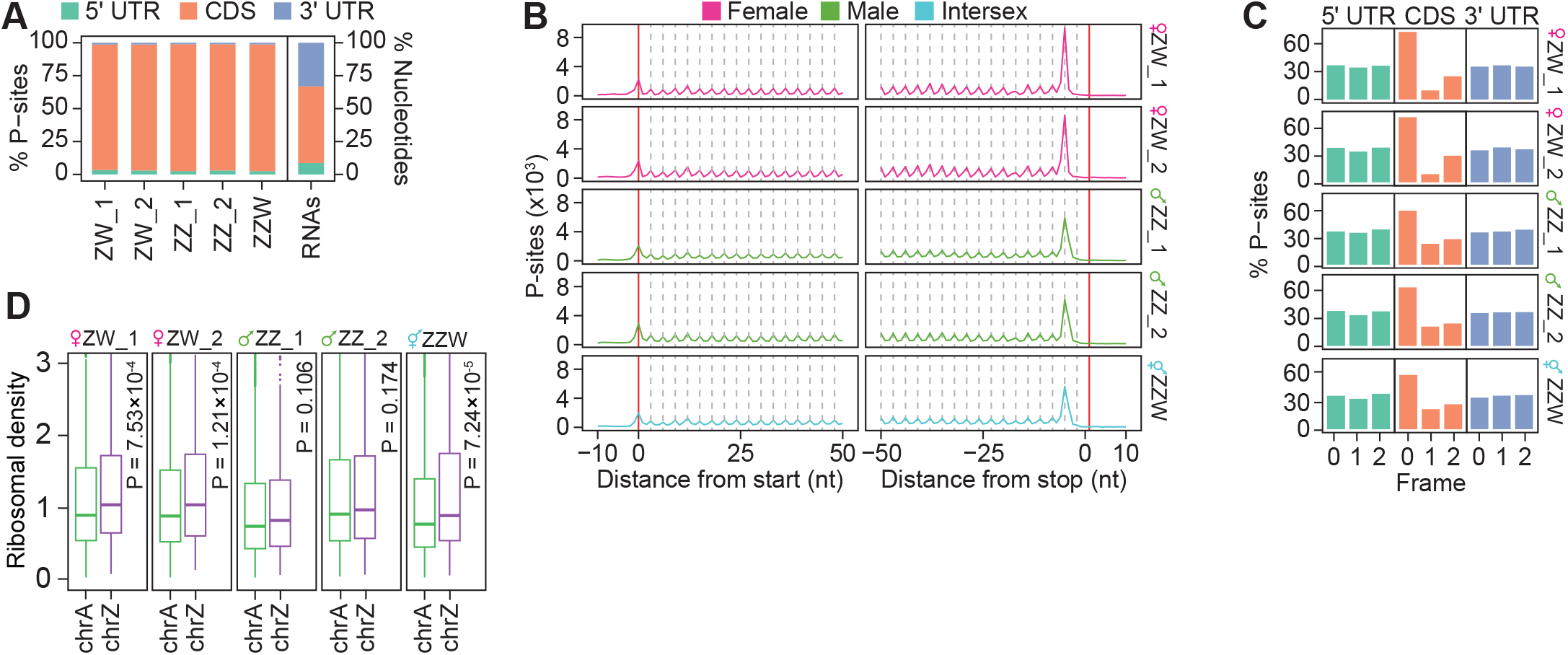
Ribosomal profiling and translational efficiency of Z-chromosome transcripts. **(A)** Bar plots displaying the percentage of ribosomal P-sites in 5’ untranslated region (UTR), coding sequence (CDS) and 3’ UTR of mRNA transcripts per sample, with the expected read distribution based on random fragmentation displayed on the right. **(B)** Metaprofile plot showing the trinucleotide periodicity along transcript CDS, with distance from start and stop codons in nucleotides on the x-axis and number of P-sites (in thousands) per position on the y-axis for each CEF line. **(C)** Bar plot representation of the percentage of ribosomal P-sites per transcript region (5’ UTR, CDS and 3’ UTR) and codon frame periodicity per CEF line. **(D)** Translation efficiency calculated as RPF (ribosome-protected fragment) counts in FPKM normalised by gene expression levels of autosomes and Z-chromosome per CEF line. Only genes with RNA FPKM > 1 and RPF FPKM > 1 were included. Mann-Whitney U-test was used for significance testing between mean translation efficiency of chrZ vs autosomes. Data shown as median, first and third quartiles and 1.5x interquartile range (IQR).

**Figure S13.**
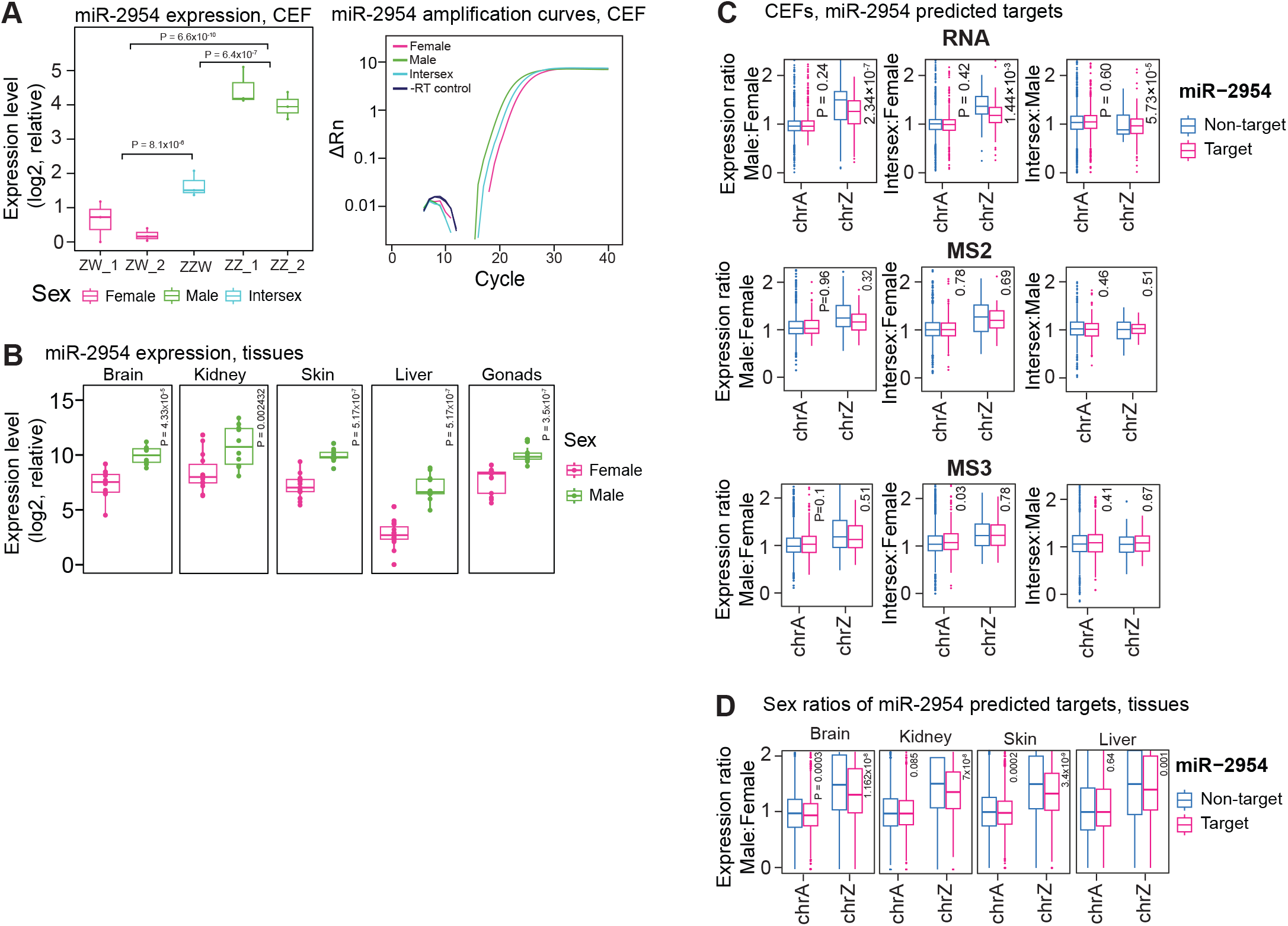
Expression of miR-2954. **(A)** miR-2954 expression in female, male and intersex fibroblasts quantified by quantitative real time PCR (qRT-PCR). Right: Boxplots of log2-relative expression of miR-2954 in female (ZW_1, ZW_2), male (ZZ_1, ZZ_2) and intersex (ZZW) fibroblasts for n=3 independent experiments. Mann-Whitney-U test used for significance testing (Males-Females, Males-Intersex, Intersex-Females). Left: Amplification curves of female, male and intersex samples, including -RT negative controls in CEFs. Females, males and intersex samples shown in pink, green and teal respectively. **(B)** miR-2954 expression in brain (n = 20, Female: n=10; Male: n=10), kidney (n=24; Female: n=14; Male: n=10), skin (n=25; Female: n=15; Male: n=11), liver (n=25; Female: n=15; Male: n=11) and gonadal tissues (n=23; Female, ovary: n=13; Male, testis: n=11) from pure (White Leghorn; WL or Red Junglefowl; RJF) and reciprocal F1 cross (Forward: RJF x WL; Reverse: WL x RJF) chicken samples. Female and male samples shown in pink and green respectively. Mann-Whitney-U used for significance testing. **(C)** Boxplots of gene expression (RNA; top panel) or protein abundance (MS2, MS3, middle and bottom panels, respectively) ratios for target (RNA: chrA=1147, chrZ=156; MS2: chrA=149, chrZ=27; MS3: chrA=236, chrZ=35) and non-target (RNA: chrA=9303, chrZ=345; MS2: chrA=1028, chrZ=27; MS3: chrA=1709, chrZ=55) genes of miR-2954 with p-values states above autosomal and Z-chromosome boxplots (Mann-Whitney-U test). Expression ratios were calculated for expressed genes (FPKM > 1 or protein abundance > 1). Data shown as median, first and third quartiles and 1.5x interquartile range (IQR). **(D)** Boxplots of Male:Female ratios of gene expression (RNA) for target (autosomal targets: n=1344-1407, Z-linked targets: n = 185-191) and non-targets (autosomal non-targets: n=10947-11843, Z-linked non-targets: n = 419-449) of miR-2954 per tissue with p-values states above autosomal and Z-chromosome boxplots (Mann-Whitney-U test). Expression ratios were calculated for expressed genes (FPKM > 1). Data shown as median, first and third quartiles and 1.5x interquartile range (IQR).

### Materials and Methods

#### Ethics statement

All animal experimental procedures were performed in accordance with Karolinska Institutet’s and Linköpings Universitet’s guidelines and approved by the Swedish Board of Agriculture (permit 16110-2020 Jordbruksverket).

#### Tissue RNA isolation

Brain, kidney, liver, skin, ovaries and testes were isolated from adult chickens. RNA isolation was performed using the phenol-chloroform extraction method. Briefly, 50-100mg of tissue was isolated and homogenised using 1 ml of TRIzol reagent [Thermofisher] and a tissue homogeniser. To precipitate the RNA, 500µl of isopropanol was added per 1 ml of TRIzol and the mixture was incubated for 10 minutes followed by centrifugation at 12000 rpm at 4°C for 10min. To wash the RNA, the pellet was resuspended in 1 ml of 75% ethanol per 1 ml of TRIzol used and centrifuged for 5 min at 7500 rpm and 4°C. The supernatant was discarded, and the pellet was left to air-dry for 5-10 min. The pellet was resuspended in 50µl of nuclease-free water, incubated at 55°C for 15 min and stored in -80°C. The concentration was measured using a Nanodrop 2000 instrument.

#### Derivation and culturing of cell lines

Chicken embryonic fibroblasts (CEFs) were derived from 10- to 13-day-old chicken embryos. Briefly, eggshells were swabbed with 70% ethanol before removing the round end of the egg. Using sterile forceps, the chorioallantoic membrane was snipped and the chicken embryos were removed. The embryos were sacrificed by decapitation and were subsequently finely minced with sterile scissors. The tissue was further dissociated by incubation in 0.25% trypsin at 37°C for 15 min. The supernatant was removed and centrifuged at 1200 rpm for 5 min and the cell pellet was washed twice in 1x PBS [Thermofisher]. The pellet was resuspended in 10 ml of complete media (10% Fetal Bovine Serum [Gibco], 100U/ml Penicillin - 100ug/ml Streptomycin, 1mM non-essential amino acids [Gibco], 1mM Sodium pyruvate [Gibco]) and plated in Petri dishes pre-coated with 0.1% gelatin solution. CEFs were incubated at 37°C in a humidified 5% CO2 incubator until confluency was reached. Upon confluency, cells were passaged using the TrypLE dissociation reagent [Thermofisher] and re-plated in 0.1% gelatin-coated plates.

#### Quantitative Real-Time PCR - miR-2954 (CEF and tissues)

##### RNA isolation

RNA isolation was performed using the phenol-chloroform RNA extraction method. Briefly, 350 µl of TRIzol reagent was added to pellets of chicken embryonic fibroblasts grown to 80% confluency, followed by addition of 70 ul of chloroform to allow for phase separation. After a room temperature incubation of 5 min, the samples were centrifuged for 15 min at 12000 g and 4°C. The aqueous phase was carefully isolated to avoid any contact with the DNA- and protein-containing inter- and organic phases and transferred to a new microcentrifuge tube. To precipitate the RNA, 175 µl of isopropanol was added and the mixture was incubated for 10 min at RT, followed by centrifugation at 12000 g and 4°C for 10 min. The supernatant was discarded, and the RNA pellet was washed with 350 µl of 75% EtOH, followed by centrifugation at 7500 g and 4°C for 5 min. Finally, the pellets were resuspended in 40 µl of RNase-free water and concentration measured on a Nanodrop 2000 instrument. *Reverse Transcription*. Reverse transcription was performed using the PrimeScript RT reagent kit with gDNA eraser [Takara] according to the manufacturer’s instructions and as previously described in Cheng et al. 2023. Briefly, 1 µg of RNA was treated with 1 µl of gDNA eraser and incubated for 2 min at 42°C, followed by reverse transcription using either Primescript’s RT primer mix at 37°C for 15 min, 85°C for 5s and 4°C hold or a miR-2954-specific stem-loop primer (RT-miR-2954-stem-loop primer: 5’-CTCAACTGGTGTCGTGGAGTCGGCAATTCAG TTGAGTGCTAGGA-3’) at 42°C for 15 min, 85°C for 5s and 4°C hold. *qPCR*. qPCR was performed using the PowerUp SYBR green mastermix reagent according to the manufacturer’s instructions. Specifically, 1 µl of 1:10-diluted cDNA was used as input in a 10 µl reaction including 5 µl SYBR green reagent, 3 µl RNase-free water and 0.5 µl forward and reverse primers at 10 µM. The reaction was performed on an Applied Biosystems Real-Time PCR instrument using the following program: 50°C for 2 minutes, 95°C for 2 minutes, followed by 40 cycles of: 95°C for 3 seconds and 60°C for 30 seconds. All Ct values are available in table S5.

#### Bulk RNA-sequencing using UMI-Smart-seq2

##### Library preparation

###### Cell lysis

Bulk Smart-seq2 was performed as previously described with slight modifications (*23*). Specifically, 2ng of purified tissue RNA was added to 3µl of Smart-seq2 lysis buffer (1uM oligo-dT primer [5’-Biosg//idSp//idSp//idSp/ACGAGCATCAGCAGCATACGAT30VN; IDT], 0.5mM (each) dNTPs, 0.2% Triton-X-100, 1 U/µl RNase inhibitor [Takara]. To ensure RNA denaturation, the samples were incubated at 72°C for 3 minutes and immediately placed on ice. *Reverse transcription and cDNA synthesis:* 5.7 µl of reverse transcription mastermix (1x Superscript II first-strand buffer, 5mM betaine [Sigma], 6mM MgCl2 [Ambion], 1µM TSO [5’-biotin-AGAGACAGATTGCGCAATGHHHHHHrG+GG-3’; IDT], 1.7U/µl of recombinant RNase inhibitor [Takara], 17U/µl Superscript II reverse transcriptase) was added to each sample and the reaction took place as follows: 42°C for 90min, 10 cycles of 50°C for 2 min, 42°C for 2 min, followed by, 70°C for 15 min and 4°C on hold. For the pre-amplification PCR, 15µl of PCR mastermix was added to each sample (1x Kapa HiFi HotStart ReadyMix [Roche], 0.1µM forward primers [5’-TCGTCGGCAGCGTCAGATGTGTATAAGAGACAGATTGCGCAATG-3’; IDT] and 0.1µM reverse primers [5’-ACGAGCATCAGCAGCATACGA-3’, IDT]) and the reaction took place as follows: 98°C for 3 min, 8 cycles of 98°C for 20 sec, 67°C for 15 sec, 72°C for 6 min, and followed by 72°C for 5 min, and 4°C on hold. *cDNA purification*. Purification of the cDNA libraries was performed by combining the cDNA samples to 22% PEG magnetic beads or AMPure XP beads at a ratio of 1:0.8. Briefly, the mixture was incubated at room temperature for 8 minutes and on the magnetic rack for 5 minutes. The supernatant was removed, and the bead pellet was washed twice with freshly prepared 80% ethanol. The bead pellet was left to air-dry for 3 minutes, and the cDNA libraries were eluted in 17µl of EB buffer [Qiagen]. Quantification of the cDNA libraries was performed using the Quantifluor dsDNA kit [Promega]. The libraries were normalised to 1ng/ul. *Tagmentation*. 2ng of cDNA was combined with 18µl of tagmentation mix containing 10mM TAPS-NaOH [Sigma], 5mM MgCl2 [Thermofisher], 8% PEG-8000 and 0.5 µl of in-house produced Tn5 at 44.5uM. The samples were incubated at 55°C for 8 minutes. To strip the Tn5 off the cDNA, 3µl of 0.2% SDS solution was added to each sample and the mixture was incubated at room temperature for 5 minutes. *PCR amplification*. 1.5µl of combined Nextera i7 and i5 [IDT] were added to each sample as well as 25 µl of PCR mastermix (1x KAPA HiFi PCR buffer, 0.06mM (each) dNTPs, 1U KAPA HiFi polymerase). The reaction took place as follows: 72°C for 3min, 95°C for 30 sec, 10 cycles of 95°C for 10 sec, 55°C for 30 sec, 72°C for 30 sec followed by 72°C for 5 min and 4°C on hold. The libraries were pooled and purified as described above. The final concentration of the pool was measured on a Qubit 4.0 using the Qubit dsDNA High Sensitivity Assay kit [Molecular probes] and the library fragment distribution was inspected on an Agilent 2100 Bioanalyzer using Agilent High Sensitivity DNA chips. The library pool was sequenced on a Nextera Nextseq 550 using a Nextseq 500/550 High-Output 75 cycle sequencing kit v2.5 [Illumina #20024906] with the following settings: Read 1 = 72 cycles, Index 1 = 10 cycles, Index 2 = 10 cycles.

#### RNA-seq data quantification

Raw multiplexed RNA-seq data was aligned and quantified to the GRCg6a genome (Gallus_gallus.GRCg6a.dna_sm.toplevel.fa.gz + Gallus_gallus.GRCg6a.100.gtf transcript annotations) using zUMIs (*49*) (v.2.9.4c, UMI(12-19), find_pattern: ATTGCGCAATG, additional_STAR_params: ‘--clip3pAdapterSeq CTGTCTCTTATACACATCT’) with barcode- and UMI filtering cutoffs allowing 1 base at phred 20 using a list of expected barcodes for edit distance-based binning.

##### Variant calling and allelic quantification

For demultiplexed and aligned bam files, read groups were added and samples were merged according to genotype using GATK (*50*) (v4.1.3.0, AddOrReplaceReadGroups, MergeSamFiles [gatk --java-options “-Xmx128G”]). Variants were called using bcftools (*51*) (v.1.10.2) mpileup (--max-depth 8000 --skip-indels) and call (-mv, in ploidy mode) then filtered for a depth over 5 reads with an allele frequency >50% using bcftools filter (-i ‘DP>5 & AF>0.5 & QUAL>10’). Next, “unique WL” variants were subsetted from RJF variants using bcftools isec (-C -w 1) and “common RJF” variants were subsetted from the “unique WL” list. As a 2nd pass filtering, allelic expression was quantified for “unique WL” variants and variant-level count tables were calculated using zUMIs with a standard GRCg6a reference genome (see below) and variants with an agreement with chicken strain in <50% of males or females were excluded to form the “final WL” variant list. Next, a custom GRCg6a reference genome was created by first inserting “common RJF” variant bases to correct for strain deviations using bcftools consensus (v.1.10.2) followed by N-masking using “final WL” variants using bcftools consensus (v.1.10.2, --mask). A STAR index was created using the WL N-masked GRCg6a genome and used for zUMIs alignment and quantification. Allelic quantification was performed on the zUMIs output bam files as previously described (*24, 36*). Briefly, variants were mapped to transcriptome positions and intersected with bases overlapping N-masked positions of the genome using the CIGAR string and reads were assigned to RJF/WL genotypes if >0.66 of basecalls matched the genotype and allelic read counts were summarised per gene and cell.

#### CEF Truseq bulk RNA-seq

##### Library preparation

Bulk RNA-sequencing using Illumina’s TruSeq RNA Library Prep Kit v2 was performed in triplicates using 500 ng of purified RNA per sample according to the manufacturer’s instructions. Briefly, the samples were incubated at 65°C for 5 min, followed by bead purification to separate and elute polyA RNA-bound beads. The eluted RNA was incubated at 94°C for 8 min followed by 4°C on hold to elute, fragment and prime the RNA for first-strand synthesis. First strand synthesis was performed by adding 8 µl of first-strand master mix (containing 1ul of Superscript II reverse transcriptase for each 9 µl First Strand Master mix) to each sample and running the following program: 25°C for 10 min, 42°C for 50 min, 70°C for 15 min and hold at 4°C. For second strand synthesis, 25 µl of second strand master mix was added to each sample, followed by incubation at 16°C for 1h. After AMPure XP bead purification, end repair was performed by adding 40 µl End Repair Mix to each sample and incubating at 30°C for 30 min. After bead purification, the 3’ ends were adenylated by adding 12.5 µl A-tailing mix to each sample followed by incubation at 37°C for 30 min, 70°C for 5 min and hold at 4°C. Indexing adapters were ligated by adding 2.5 µl Ligation mix and 2.5 µl RNA adapter Index (unique to each sample) per sample and the mixture was incubated at 30°C for 10 min followed by addition of 5 µl of Stop Ligation buffer to each sample to stop the ligation reaction. After bead purification, DNA fragments were enriched through PCR by adding 5 µl PCR primer cocktail and 25 µl PCR Master Mix to each sample. The reaction was performed using the following program: 98°C for 30s, 15 cycles of 98°C for 10s, 60°C for 30s, 72°C for 30s followed by 72°C for 5 min and on hold at 10°C. Following a final AMPure XP bead purification, the library quality and size was validated by running the samples on a High Sensitivity dsDNA Bioanalyzer chip and quantified by real-time quantitative PCR (RT-qPCR). Libraries were pooled in equimolar amounts and sequenced on a Nextseq 550 instrument using a Nextseq 500/550 High-Output 75 cycle sequencing kit v2.5 [Illumina #20024906] with the following settings: Read 1 = 76 cycles, Index 1 = 6 cycles, Index 2 = 6 cycles.

##### Data analysis

Raw BCL files were converted to FASTQ and demultiplexed using bcl2fastq (v.2.20.0.422). The demultiplexed raw data was aligned and quantified to an N-masked GRCg6a genome (see section *Variant Calling and allele quantification* under *Bulk RNA-sequencing using UMI-Smart-seq2*) and transcriptome (Gallus_gallus.GRCg6a.100.gtf transcript annotations) using zUMIs (v.2.9.4c, cDNA (1-76), BC(1-8), additional_STAR_params: ‘--clip3pAdapterSeq AGATCGGAAGAGCACACGTCTGAACTCCAGTCA’) with barcode filtering cutoffs allowing 1 base at phred 20 using a list of expected barcodes for edit distance-based binning. Allele calling was performed as described in section *Variant Calling and allele quantification* under *Bulk RNA-sequencing using UMI-Smart-seq2*.

#### Single-cell RNA-sequencing using Smart-seq3

##### Library preparation

All scRNA-seq libraries were prepared as previously described (*29*). Briefly, chicken embryonic fibroblasts (CEFs) were sorted into 384-well PCR plates [Thermofisher] containing 3µl of lysis buffer (5% PEG-8000 [Sigma], 0.1% Triton-X-100 [Sigma], 0.5 units/µl RNase Inhibitor [Takara], 0.5mM (each) dNTPs [Thermofisher], 1uM oligo-dT primer [5’-Biotin-ACGAGCATCAGCAGCATACGAT30VN-3’; IDT]. Sorting was performed using a FACS Aria II. After sorting, the plates were sealed, briefly centrifuged, and stored in -80°C. To ensure cell lysis and RNA denaturation, the plates were incubated at 72°C for 10 min and immediately placed on ice. For reverse transcription, 1µl of reverse transcription master mix (25mM Tris-HCl pH 8.3 [Sigma], 30mM NaCl [Ambion; Thermofisher], 2.5mM MgCl2 [Ambion; Thermofisher], 1mM GTP [Thermofisher], 8mM DTT [Thermofisher], 0.5 units/µl RNAse Inhibitor [Takara], 2uM template-switching oligo [5’-biotin-AGAGACAGATTGCGCAATGNNNNNNNNrGrGrG-3’; IDT], 2U/µl Maxima H-RT enzyme [Thermofisher]) was added to each sample. Reverse transcription was performed at 42°C for 90min, followed by 10 cycles of 50°C for 2min and 42°C for 2min, and terminated at 85°C for 5min. For PCR pre-amplification, 6µl of PCR master mix (1x KAPA HiFi HotStart Buffer [Roche], 0.3mM (each) dNTPs [Thermofisher], 0.5mM MgCl2 [Thermofisher], 0.5uM forward primer [5’-TCGTCGGCAGCGTCAGATGTGTATAAGAGACAGATTGCGCAA-3’; IDT], 0.1uM reverse primer [5’-ACGAGCATCAGCAGCATAC*G*A-3’; IDT], 0.02U/µl polymerase) was added to each sample. PCR pre-amplification was performed using the following thermocycler program: 98°C for 3min, 20 cycles of 98°C for 20 sec, 65°C for 30 sec, 72°C for 4 min, followed by 72°C for 5 min and 4°C on hold. cDNA purification was performed using in-house prepared 22% PEG beads at a beads-to-sample ratio of 0.6:1. cDNA was quantified using the Quantifluor dsDNA kit [Promega]. cDNA was normalised to a final concentration of 100pg/µl. For the tagmentation step, 100pg of cDNA were incubated with 1 µl of tagmentation mastermix (0.1µl of tagmentation buffer 4x containing 40mM Tris-HCl pH 7.5, 20mM MgCl2, 20% Dimethylformamide, 0.1µl Amplicon Tagment Mix -Tn5 [Nextera], 0.40µl water) at 55°C for 10 min. To strip the Tn5 from the cDNA, 0.5 µl of freshly prepared 0.2% SDS solution [Sigma] was added to each sample and incubated at room temperature for 5 min. The samples were indexed using 1 µl of 1uM in-house, pre-mixed Nextera index primers [IDT] and post-tagmentation PCR was performed by adding 3µl of PCR mastermix (1.4 µl Phusion HF 5x buffer, 0.2mM (each) dNTPs, 0.01U/µl Phusion HF polymerase) to each sample. PCR was performed using the following program: 72°C for 3min, 98°C for 3min, 10 cycles of 98°C for 10 sec, 55°C for 30 sec, 72°C for 30 sec, followed by 72°C for 5 min and 4°C on hold. The samples were subsequently pooled and purified using in-house 22% PEG magnetic beads with a ratio of beads-to-sample of 0.7:1.

#### Single-cell RNA-seq using UMI spike-ins

##### Library preparation

Full-length single-cell RNA-seq library preparation using the Xpress-seq (v1) method was performed at Xpress Genomics (Stockholm, Sweden). In brief, single cells were sorted using a Sony SH800S instrument into provided 384-well plates containing lysis buffer, spun down and stored at -80 °C. Upon submitting plates to Xpress Genomics, robotic automated library preparation was performed. Sequencing was performed on the DNBSEQ G400RS platform (MGI Tech) using App-C Sequencing primers.

##### Data analysis

Data was pre-processed using zUMIs (v.2.9.7) as described above but with the following modifications: 2 mismatches were allowed in detection of UMI-read patterns, and for barcode and UMIs, 4 and 3 mismatches were allowed, respectively. Additionally, spike-in sequences for the 5’ complex set of molecular spikes were included as mappable sequences (https://raw.githubusercontent.com/sandberg-lab/molecularSpikes/main/fasta_reference/molecularSpikes_complexset_5p.fa). Molecular spikes were extracted from aligned bam files using the UMIcountR package(*33*) (https://github.com/cziegenhain/UMIcountR) and overrepresented spike-ins were removed (>5 barcode or >100 sequences). Next, cells with less than 10% reads in spike-ins were kept and outliers were detected based on low gene detection (log 3 MADs) or read counts (log 5 MADs) and excluded. Spike-in size factors were calculated and UMIs were normalised using scater/scran (v.1.24.0, computeSpikeFactors, logNormCounts transform = “none”).

#### Public DNA methylation data analysis

Raw whole-genome bisulfite sequencing data for white leghorn samples was obtained from SRR1258373, SRR1258374, SRR1258375 and SRR1258376. Data was quality- and adapter trimmed and low complexity reads were excluded using fastp (--low_complexity_filter –detect_adapter_for_pe) and aligned to the GRCg6a genome using abysmal (*52*) (v.3.2.2, default settings). Sex was determined by calculating read coverage of chromosome W. 5mC methylation levels was calculated for CpG sites only and symmetrical CpG methylation counts was summarised using dnmtools (*53*) (v.1.4.2, format, counts -cpg-only, sym). Methylation counts were merged per sex using dnmtools merge and summarised using dnmtools merge (-t). CpGs with at least 5 total read counts in both males and females were kept. For genome-wide visualisation, methylated- and total counts were aggregated into 10kb bins where bins within 1Mb of centromeres (obtained from the UCSC table browser gap track) were excluded. Methylation fractions were calculated as methylated counts / total counts.

#### Genome-wide mappability calculation

The GRCg6a genome (Gallus_gallus.GRCg6a.dna_sm.toplevel.fa) was indexed and k50-mer mappability was calculated using genmap (*54*) (v.1.3.0, map -K50 -E 0). Mappability was rounded to two decimals using awk (v.5.1.0, ‘{OFS = “\t”;$4=sprintf(“%.2f”,$4)}1’), coordinate-sorted then converted to bigwig format using the UCSC tool bedGraphToBigWig (v.4). To identify contiguous regions of low mappability, sliding windows were created across the genome using bedtools (v.2.30.0, makewindows -w 5000 -s 1000) and mappability per window was calculated using deeptools (v.3.5.4.post1, multiBigwigSummary BED-file). Next, windows with <80% mappability were extracted using awk and windows within 10kb distance were merged using bedtools (v.2.30.0, merge -d 10000). Regions of >10kb were kept and used for masking low mappability regions (called ‘lowmap’ below) where indicated.

#### DNA-sequencing

##### Library preparation

Genomic DNA was isolated from cultured CEFs using the Monarch Genomic DNA purification kit according to the manufacturer’s instructions. The concentration of gDNA was quantified using a Nanodrop 2000 instrument and the gDNA was subsequently diluted to 1ng/ul. For DNA-sequencing, gDNA tagmentation was performed using an in-house prepared Tn5 enzyme as previously described (*55*). Briefly, 5ng of gDNA was incubated with 15µl of tagmentation mastermix (10mM TAPS [Sigma], 5mM MgCl2 [Thermofisher], 10% Dimethylformamide [Sigma], 2.25uM Tn5 [in-house]) at 55°C for 8 minutes. To strip the Tn5 from the DNA, 3.5µl of 0.2% SDS was added to each reaction. The samples were quickly centrifuged and incubated at room temperature for 5 min. The samples were indexed using 2.5µl of 1 µM pre-mixed Nextera index primers [IDT]. Post-tagmentation PCR was performed by adding 16.5µl of PCR master mix (1x KAPA HiFi PCR buffer, 0.6mM (each) dNTPs, 1U/µl KAPA HiFi polymerase) to each sample and incubating using the following program: 72°C for 3min, 95°C for 30sec, 6 cycles of 95°C for 10 sec, 55°C for 30sec, 72°C for 30sec, followed by 72°C for 5 min and 4°C on hold. Double purification of the final libraries was performed using in-house 22% PEG magnetic beads (*29*). Briefly, 22% PEG magnetic beads were combined with the pooled DNA libraries in a bead-to-sample ratio of 0.9:1 and incubated at room temperature for 8 minutes. The samples were then placed on a magnetic rack for 5 minutes. The clear supernatant was then removed and discarded, and the bead pellets were washed twice with freshly prepared 80% EtOH. The beads were left to air-dry for 3 minutes while remaining on the magnetic rack. The samples were eluted in 30 µl. To ensure complete removal of residual impurities and primer-dimers, the purification was repeated as described above and the final sample was eluted in 17 µl of nuclease-free water [Ambion]. Library fragment size was assessed using a Bioanalyzer high-sensitivity dsDNA chip and library concentrations were quantified using Qubit’s high-sensitivity dsDNA quantification kit on a Qubit 3.0 Fluorometer. Libraries were pooled in equimolar amounts and sequenced on a Nextseq 550 instrument using a Nextseq 500/550 High-Output 75 cycle sequencing kit v2.5 [Illumina #20024906] with the following settings: Read 1 = 74 cycles, Read 2 = 74 cycles, Index 1 = 10 cycles, Index 2 = 10 cycles.

#### DNA-seq data analysis

Raw DNA-seq data was adapter- and quality trimmed using fastp (*56*) (v.0.20.0, –adapter_sequence CTGTCTCTTATACACATCT –adapter_sequence_r2 CTGTCTCTTATACACATCT) and aligned to the GRCg6a reference genome using minimap2 (*57*) (v.2.24-r1122, -ax sr). Reads were sorted, mate-pair information fixed, and duplicates marked using biobambam2 (v.2.0.87, bamsort fixmates=1 markduplicates=1). Variants were called using bcftools mpileup (–ignore-RG -a AD,DP, –max.depth 8000) and call (-mv, in ploidy mode) using ZZ and ZW ploidies with sample-sex information. Variants within 5bp of indels were excluded and heterozygous variants sequenced to a depth over 5 reads with a minor allele frequency over 10% were filtered using bcftools filter (-g 5 -i ‘TYPE=“snp” & QUAL >10 & INFO/DP>5 & GT=“het” & MAF>0.1’). To calculate DNA copy numbers, the GRCg6a genome was binned into 100kb bins using bedtools (*58*) makewindows (v.2.30.0, -w 100000) and binned read counts were calculated using bedtools multicov (-q 13). Genome statistics were also calculated for the same bins; mappability (see above) using deeptools (multiBigwigSummary BED-file); nucleotide frequencies using bedtools (nuc); assembly gaps were obtained from UCSC and gaps >1kb were kept and bins within 500kb were identified using bedtools (window -w 500000 -c); RepeatMasker rmsk track was obtained from UCSC and overlapped with bins using bedtools (intersect -wao | map -c 10) and percentage overlap was calculated. Bins with >2.5% N bases or average mappability <50% or within 500kb of a large assembly gap or with a rmsk fraction 2 MADs above median were excluded. Data was corrected for GC-content and mappability and DNA copies were estimated using HMMcopy (v.1.38.0, correctReadcount mappability = 0.8). Expected ploidies were set to 2 for diploid samples and 3 for triploid and multiplied with DNA copies to adjust for ploidy and regions annotated as ideal by HMMcopy were used for plotting. For base-resolution variants, only variants with a read depth of 6-50, heterozygous genotype and >0 variance were kept and variants overlapping excluded genome bins were removed.

#### ATAC-seq

##### Library preparation

Omni ATAC-seq libraries were prepared as previously described (*59*) with slight modifications. Briefly, 100 000 cells were pelleted at 500 RCF at 4°C for 5 min. The supernatant was carefully removed, and the pellet was resuspended in 50µl ATAC-RSB lysis buffer (10mM Tris-HCl pH 7.4, 10mM NaCl, 3mM MgCl2, 0.1% NP-40, 0.1% Tween-20, 0.01% Digitonin) and incubated on ice for 10 min. The lysis buffer was washed out by adding 1ml of ATAC-RSB wash buffer containing 0.1% Tween but no NP-40 or digitonin and the samples inverted 3 times to mix, and the nuclei pelleted at 500 RCF at 4°C for 10 min. The supernatant was removed, and the samples were resuspended in 50µl transposition mixture (25µl 2x TD buffer, 1.5µl Tn5 (27µM) 16.5 µl 1x PBS, 0.5 µl 1% digitonin, 0.5 µl 10% Tween-20, 7µl water). Tagmentation was performed in a thermoshaker at 37°C for 30 min at 1000rpm. The transposed DNA was purified using the Zymo DNA Clean and Concentrator-5 kit following the manufacturer’s instructions and eluted in 21µl of elution buffer. Library amplification was performed by adding 30 µl of PCR master mix to the purified DNA (25µl 2x NEBNext High Fidelity PCR Master Mix, 2.5 µl Ad1_noMX (common i5 Nextera adapter primer), 2.5 µl Ad2 (unique i7 Nextera adapter primer). The libraries were amplified for 11 cycles using the following cycling conditions: 72°C for 5 min, 98°C for 30s, 11x (98°C for 10s, 63°C for 30s, 72°C for 1 min), 4°C on hold. The final libraries were fragment size-selected by double-sided 0.5x/1.3x bead purification using homemade 22% PEG magnetic beads. Briefly, 25 µl of room temperature Ampure XP beads were added to each sample (beads-to-sample ratio = 0.5) and incubated for 10 min after thorough resuspension. The samples were placed on a magnetic rack and the supernatant was removed and transferred to a new microcentrifuge tube containing 65 µl room temperature Ampure XP beads (beads-to-sample (original volume) ratio = 1.3). After thorough mixing, the samples were incubated at room temperature for 10 min and placed on a magnetic rack for 5 min. The supernatant was discarded, and the beads were washed twice with 200µl of freshly prepared 80% ethanol. After ethanol removal, the samples were air-dried for 5 min and the libraries eluted in 20µl of nuclease-free water. The libraries were pooled in equimolar amounts and paired-end sequencing was performed on an Illumina Nextseq 550 instrument to obtain ∼ 20 million paired-end reads per sample.

##### Data preprocessing

Raw ATAC-seq data was converted to FASTQ format using bcl2fastq (v.2.20.0.422). Sequencing adapter removal was performed using fastp (v.0.20.0). The resulting trimmed reads were aligned to the GRCg6a genome using Bowtie 2 (v.2.5.1, settings: -N 1, -X 2000, --very-sensitive). Aligned reads were then converted to BAM format, mate pair information was fixed, duplicate read pairs were marked, and the BAM output was sorted according to read coordinates using biobambam2 (v.2.0.87, bamsort fixmates=1, markduplicates=1). Replicate merging was performed using sambamba merge (v. 0.7.0). SNPs were called from the ATAC-seq data using bcftools (v. 1.10.2, mpileup --ignore-RG -Ou -a AD,DP --max-depth 8000 | call --threads 64 -mv -Oz --ploidy-file –samples-file | filter -g 5 -i ‘TYPE=“snp” & QUAL>10 & INFO/DP>5 & GT=“het” & MAF>0.1’). Next, the GRCg6a genome was N-masked for SNP positions using bcftools (consensus --mask) and data was realigned to the N-masked reference using bowtie2 as described above. Peak calling on all replicates was performed using Genrich in ATAC-seq mode (v.0.6, settings: -j, -r, -d 150, -e W,MT, -E lowmap), PCR duplicates were removed, the cut sites were expanded to 150 bp and reads from the W chromosome and the mitochondria were excluded, and it was repeated both with and without low mappability regions. Genrich requires input sorted by query name which was done by biobambam2 bamsort. All peaks were combined and replicated peaks within 100bp were merged and counted using bedtools (v.2.30.0, merge -c 1 -o count -d 100) and peaks present with >1 counts were kept. Distance to nearest TSS for each peak was annotated using bedtools (v.2.30.0, closest). Peak quantification of proper read pairs, both excluding and including low-mapping regions, and removing PCR duplicates was calculated by deepTools (v.3.5.4.post1, multiBamSummary -- samFlagInclude 2, --ignoreDuplicates, -bl lowmap). Genome-wide signal pileups was normalised using deepTools (bamCoverage --normalizeUsing RPGC, --effectiveGenomeSize 1058535536, -- ignoreDuplicates, --samFlagInclude 2, -ignore MT, --minFragmentLength 38, --maxFragmentLength 2000). These only included proper read pairs, mitochondrial entries were removed as well as PCR duplicates, and only fragments between 38 and 2000 bp lengths were kept. The pileups were also made by containing, and excluding, low mapping regions, for further use and visualisations. Effective genome size of GRCg6a used for normalisation was obtained from Genrich. Enrichment 5kb around TSS was calculated using deeptools (computeMatrix -a 5000, -b 5000 -R Gallus_gallus.GRCg6a.100.gtf).

##### Allelic analysis

To phase the Z chromosome, all SNP positions were extracted using bcftools corresponding to all SNPs (‘all_snps’; query -f ‘%ID\t%CHROM\t%POS\t1\t%REF/%ALT\n’) and chrZ SNPs with a genotype matching the non-present allele in either female sample (‘female_gt_snps’; query -i ‘CHROM = “Z” && (GT[0] = “0” | GT[1] = “1”) ‘ -f ‘%ID\t%CHROM\t%POS\t1\t%ALT/%REF\n’). Next, REF and ALT bases were flipped for SNPs not matching genotypes in females using awk (‘NR==FNR{a[$2,$3]=$5;next}(($2,$3) in a){OFS=“\t”;$5=a[$2,$3]}1’ female_gt_snps all_snps). These corrected SNPs were used as input to SNPsplit (v.0.4.0, –no_sort –snp_file corrected_snps) to split the data into respective alleles. Peak calling, peak quantification and TSS enrichment were performed as described above for each allele.

##### Transcription factor footprinting and identification of differentially bound transcription factors

Differential footprinting analysis was performed on called peaks (see section ATAC-seq *data analysis*) using TOBIAS v0.16.1 (default settings). ATACorrect was applied to correct for Tn5 bias and resolve hidden footprints. To identify regions of protein binding across the genome, footprinting scores were calculated across the open chromatin regions, using the corrected signals, with the TOBIAS ScoreBigwig. Differential TF binding analysis was performed with TOBIAS BINDetect module which combines footprinting scores with TF motif information from JASPAR CORE 2024 (non-redundant, vertebrate). The heatmap was generated using ComplexHeatmap and the motif clustering using motifStack.

#### Proteomics

##### Sample preparation

Cell pellets of approximately 1 million cells were collected at 300g for 5 min and washed with ice-cold PBS 5 times to eliminate serum-containing media. Cell pellets were solubilized in 20 µl of 8M urea in 50 mM Tris-HCl, pH 8.5 sonicated in water bath for 5 min before 10 µl of 1% ProteaseMAX surfactant (Promega) in 10% acetonitrile (ACN) and Tris-HCl as well as 1 µl of 100x protease inhibitor cocktail (Roche) was added. The samples were then sonicated using VibraCell probe (Sonics & Materials, Inc.) for 40 s with pulse 2-2 s (on/off) at 20% amplitude. Protein concentration was determined by BCA assay (Pierce) and a volume corresponding to 25 μg of protein of each sample was taken and supplemented with Tris-HCl buffer up to 90 µl. Proteins were reduced with 3.5 µl of 250 mM dithiothreitol in Tris-HCl buffer, incubated at 37°C during 45 min and then alkylated with 5 µl of 500 mM iodoroacetamide at room temperature (RT) in dark for 30 min. Then 0.5 μg of sequencing grade modified trypsin (Promega) was added to the samples and incubated for 16 h at 37°C. The digestion was stopped with 5 µl cc. formic acid (FA), incubating the solutions at RT for 5 min. The sample was cleaned on a C18 Hypersep plate with 40 µl bed volume (Thermo Fisher Scientific), dried using a vacuum concentrator (Eppendorf). Peptides, equivalent of 25 μg protein, were dissolved in 70 µl of 50 mM triethylammonium bicarbonate (TEAB), pH 7.1 and labelled with TMTpro mass tag reagent kit (Thermo Fisher Scientific) adding 100 μg reagent in 30 µl anhydrous ACN in a scrambled order and incubated at RT for 2 h. The reaction was stopped by addition of hydroxylamine to a concentration of 0.5% and incubation at RT for 15 min before samples were combined and cleaned on a C-18 HyperSep plate with 40 µl bed volume. The combined TMT-labelled biological replicates were fractionated by high-pH reversed-phase after dissolving in 50 µl of 20 mM ammonium hydroxide and were loaded onto an Acquity bridged ethyl hybrid C18 UPLC column (2.1 mm inner diameter x 150 mm, 1.7 µm particle size, Waters), and profiled with a linear gradient of 5–60% 20 mM ammonium hydroxide in ACN (pH 9.0) over 48 min, at a flow rate of 200 µl /min. The chromatographic performance was monitored with a UV detector (Ultimate 3000 UPLC, Thermo Scientific) at 214 nm. Fractions were collected at 30 s intervals into a 96-well plate and combined in 12 samples concatenating 8-8 fractions representing peak peptide elution.

##### Liquid Chromatography-Tandem Mass Spectrometry Data Acquisition

The peptide fractions in solvent A (0.1% FA in 2% ACN) were separated on a 50 cm long EASY-Spray C18 column (Thermo Fisher Scientific) connected to an Ultimate 3000 nano-HPLC (ThermoFisher Scientific) using a gradient from 2-26% of solvent B (98% AcN, 0.1% FA) in 90 min and up to 95% of solvent B in 5 min at a flow rate of 300 nL/min. Mass spectra were acquired on a Orbitrap Fusion Lumos tribrid mass spectrometer (Thermo Fisher Scientific) ranging from m/z 375 to 1500 at a resolution of R=120,000 (at m/z 200) targeting 4×105 ions for maximum injection time of 50 ms, followed by data-dependent higher-energy collisional dissociation (HCD) fragmentations of precursor ions with a charge state 2+ to 6+, using 45 s dynamic exclusion. The tandem mass spectra of the top precursor ions were acquired in 3 s cycle time with a resolution of R=50,000, targeting 1×105 ions for maximum injection time of 150 ms, setting quadrupole isolation width to 0.7 Th and normalized collision energy to 35%.

##### Data Analysis

Acquired raw data files were analyzed using Proteome Discoverer v3.0 (Thermo Fisher Scientific) with MS Amanda v2.0 search engine against Gallus gallus protein database (UniProt). A maximum of two missed cleavage sites were allowed for full tryptic digestion, while setting the precursor and the fragment ion mass tolerance to 10 ppm and 0.02 Da, respectively. Carbamidomethylation of cysteine was specified as a fixed modification. Oxidation on methionine, deamidation of asparagine and glutamine, as well as acetylation of N-termini and TMTpro were set as dynamic modifications. Initial search results were filtered with 1% FDR using the Percolator node in Proteome Discoverer. Quantification was based on the reporter ion intensities

#### Ribosomal profiling

##### Library preparation

###### Isolation of ribosome-protected fragments (RPFs)

Cells were grown to 80 % confluency on 2x 15-cm dishes. Medium was discarded and plates were shortly submerged in liquid nitrogen to snap freeze cells. 300 µl of 2x lysis buffer (50 mM Tris pH 7.5, 200 mM NaCl, 20 mM MgCl_2_, 2 mM DTT, 200 μg/ml cyclohexamide, 2 % Triton X-100, 2x Complete EDTA-free protease inhibitor cocktail (Roche), 4000 U/ml TURBO DNase I (Thermo Fisher)) was added dropwise on each plate and lysates were collected using cell scrapers. Cell debris were removed by centrifugation (10,000 x g, 15 min, 4 °C). RNA concentrations were measured by Qubit RNA Broad Range kit (Invitrogen) and 90 μg were subjected to RNase treatment for 45 min at 22 °C (750 U, Ambion RNase I, Thermo Fisher). RNase treatment was stopped by the addition of 15 µl RNase inhibitor (1 U/µl, SUPERase-In, Thermo Fisher), which was followed by a short centrifugation step to remove insoluble material (5,000 x g, 5 min). Supernatants were loaded on 1 M sucrose cushions (25 mM Tris pH 7.5, 100 mM NaCl, 10 mM MgCl_2_, 1 mM DTT, 100 μg/ml cyclohexamide, 1x Complete EDTA-free protease inhibitor cocktail (Roche), 40 U/ml RNase inhibitor (SUPERase-In, Thermo Fisher) in 11 × 34 mm tubes (Beckman Coulter) and ribosomes were pelleted at 55.000 rpm for 3 h in a TLS-55 rotor (Beckman Coulter). Afterwards, supernatants were removed, and ribosomal pellets were resuspended in 1 ml TRIZOL reagent (Thermo Fisher). RNA was isolated according to manufacturer’s instructions. Isolated RNA was heated at 80 °C for 3 min, put on ice for 1 min, mixed with Gel Loading Buffer II (ThermoFisher) and loaded onto a 15 % Novex TBE-Urea gel (ThermoFisher). The gel was run in 1x TBE buffer at 100 V for ∼2 h. After completion of the run the gel was stained with 1x SYBR Gold Nucleic Acid Gel Stain (ThermoFisher) in 1x TBE. Nucleic acids were visualized and bands referring from 25-35 nt were excised. RNA was extracted from gel slices in 600 µl RNA extraction buffer (300 mM NaOAc pH 5.5, 1 mM EDTA, 0.25 % SDS) rotating at 4 °C overnight. The next day, RNA was precipitated by adding 1.8 ml ice-cold EtOH together with 4 µl GlycoBlue Coprecipitant (ThermoFisher) and subsequent storage at -80 °C ON. Precipitated RNA was pelleted by centrifugation (5,000 x g, 10 min, 4 °C). Pellet was once washed with 1 ml EtOH, dried for ∼5 min and resuspended in 15 µl 10 mM Tris pH 7.5 supplemented with 1 µl RNase inhibitor (SUPERase-In, ThermoFisher).

##### Ligation of adaptors to RPFs

Samples were heated at 80 °C for 2 min before placing on ice. Next, 3’ phosphates were removed by T4 PNK treatment (1 µl T4 PNK (NEB) added) in 1x T4 PNK buffer (NEB) at 37 °C for 2 h. Reaction was stopped by heat inactivation (65 °C, 10 min). RNA was pelleted by addition of 70 µl water, 2 µl GlycoBlue Coprecipitant (ThermoFisher), 10 µl 1 M NaOAc and 300 µl EtOH and subsequent storage at -80 °C. RNA was washed and dried as described earlier and finally resuspended in 7 µl 10 mM Tris pH 7.5 supplemented with 1 µl RNase inhibitor. RNA libraries were generated using TruSeq Small RNA Library Prep Kit (Illumina) according to the manufacturer’s protocol with some modifications. Preparation was started by adding 1.2 µl adenylated RA3 to dephosphorylated RNA and incubating the mixture at 80 °C for 2 min. Afterwards, ligation was performed by addition of 2 µl of T4 RNA Ligase 2 (truncated K227Q), 2 µl T4 RNA Ligase 2 buffer and 6 µl PEG8000 (all components from NEB) and incubation at 14 °C ON. RNA was precipitated as described earlier, 20 µl 3 M NaOAc and 600 µl EtOH) and resuspended in 4 µl 10 mM Tris pH 7.5. Ligation products were then purified on a 15 % Novex TBE-Urea gel (ThermoFisher), extracted, and precipitated as described earlier. Next, RNA was resuspended in 13 µl 10 mM Tris pH 7.5 supplemented with 1 µl RNase inhibitor. Then, 2 mM ATP, 2 µl 10x T4 PNK buffer and 2 µl T4 PNK (NEB) were added, and the reaction mixture was incubated for 2 h at 37 °C, followed by heat inactivation (65 °C, 10 min). RNA was precipitated and resuspended in 13 µl 10 mM Tris pH 7.5 supplemented with 1 µl RNase inhibitor. Thereafter, RNA footprints were ligated with 5’ RNA adaptor (RA5, Illumina) by adding 1.2 µl RA5, 2 µl 10x T4 buffer and 2 µl T4 RNA ligase (Promega) and incubating at 14 °C ON. RNA was precipitated and resuspended in 3 µl 10 mM Tris pH 7.5. *Reverse transcription and PCR amplification of library*. Reverse transcription was performed using RNA RT primers from TrueSeq Small RNA Library Prep Kit (Illumina) and SuperScript III First-Strand Synthesis System (ThermoFisher) according to the manufacturer’s protocol.

Afterwards, 2 µl of RT products were PCR amplified using Phusion High-Fidelity PCR master mix (NEB) and DNA primers from TrueSeq Small RNA Library Prep Kit (Illumina). The PCR products were resolved on a 10 % Novex non-denaturing TBE gel (ThermoFisher) using 1x TBE running buffer. PCR products were excised and extracted using DNA extraction buffer (300 mM NaCl, 10 mM Tris pH 8, 1 mM EDTA). Subsequently, PCR products were precipitated and pelleted. Libraries were resuspended in 12 µl 10 mM Tris pH 7.5. *Duplex-specific nuclease (DSN) digestion*. To reduce the amount of ribosomal RNA contamination DSN digestion was performed using a DSN kit (evrogen). First, 4 µl of hybridization buffer (200 mM HEPES pH 7.5, 2 M NaCl) was added to the libraries. Next, libraries were heated for 2 min at 98 °C followed by incubation for 5 h at 68 °C. Consecutively, 1x master buffer (evrogen) together with 2 µl DSN enzyme were added to the samples and incubated additional 25 min at 68 °C. Digestion was stopped by addition of 20 µl stop solution (evrogen) and 5 min incubation at 68 °C. Finally, samples were cooled down on ice and DNA was isolated by phenol/chloroform extraction. Therefore, samples were mixed with 160 µl water and 200 µl phenol/chloroform (1:1) and the aqueous phase was precipitated as before. 2 µl of digested libraries were subjected to another round of PCR amplification and consecutive gel purification. Final libraries were resuspended in 11 µl 10 mM Tris pH 7.5. To assess fragment size distribution and final library concentrations, libraries were run on a Bioanalyzer instrument using a High Sensitivity dsDNA kit and quantified using the Qubit 1x dsDNA high sensitivity kit. Libraries were then pooled in equimolar amounts. Paired-end sequencing was performed on a Nextseq550 instrument to obtain ∼20 million reads per sample using the following settings: Read 1 = 75 cycles, Index 1 = 6 cycles, Read 2 = 75 cycles.

##### Data analysis

Raw Ribo-seq BCL data was converted to FASTQ and demultiplexed using bcl2fastq (v. 2.20.0.422) followed by adapter trimming using FASTP. For pre-alignment to ribosomal RNA (rRNA) sequences, a fasta file of rRNA sequences obtained from SILVA (release 138, smr_v4.3_default_db) was indexed using bowtie2-build. Pre-alignment to rRNA sequences was performed using bowtie2 (v. 2.5.1, settings: -N 1, --very-sensitive, --al-conc, --un-conc). Unaligned bowtie2 output aligned to the GRCg6a genome and transcriptome (v.2.7.2a, genome: 2.7.2a, transcriptome: Gallus_gallus.GRCg6a.100.gtf using STAR (v.2.7.2a, --runMode alignReads, --sjdbOverhang 31, --seedSearchStarLmax 10, -- outFilterMultimapNmax 2, --quantMode TranscriptomeSAM). To ensure comparability between samples, 3.5 million reads were downsampled for each sample prior to P-site assignment and downstream analyses.

Downstream analysis including P-site assignment and transcript-level quantification was performed using Ribowaltz (v.2.0). PCR duplicates were removed with the option duplicates_filter (extremity = “both”) and p-site offsets were calculated with default settings (flanking=6, extremity = “auto”). The per-gene sums of ribosome fragment counts mapping to the coding sequence (CDS) of protein-coding transcripts were used to calculate FPKM-normalised Ribo-seq counts, which were used in downstream analyses. For all paired Ribo-seq/RNA-seq analyses, bulk RNA-seq data (Truseq) was used, the translation index was calculated as Ribo-FPKM/RNA-FPKM per gene and sample. Z:autosome ratios were calculated relative to autosomal translation indexes. Similarly, Female:Male ratios were calculated as the mean translation index per gene and chromosome. Additionally, a bootstrapping method was used to correct for the difference in gene content between autosomes and the Z chromosome and has been applied both for calculations of Female:Male ratios and Z:autosome ratios of translational efficiency.

#### Multiplexed Quantitative ChIP-seq

##### Library preparation

Triplicate pellets of 10^6^ cells were collected for all conditions, flash frozen and stored at −80 °C before use. Immunoprecipitation was performed using the EpiFinder Genome kit (Epigenica, EpGe001) according to the manufacturer’s instructions.

In brief, native frozen cell pellets were lysed and MNase digested to mono- to tri-nucleosome fragments and ligated with double-stranded DNA adaptors in a one-pot reaction. Barcoded samples were then pooled and aliquoted into individual ChIP reactions with Protein A (Dynabeads; Thermofisher) for the following antibodies: H3K4me3 [Millipore; 04-745], H3K27ac [Active Motif; 39034], H3K9ac [Active Motif; 39137-AF], H4K16ac [Millipore; 07-329]. Briefly, 260 µl of sample pool was treated with 1 µl RNase A and 3 µl Proteinase K, followed by digestion at 37°C for 15 min and at 63°C for 45 min with agitation at 1000 rpm. AMPure XP bead purification was performed at 1:1 ratio of sample:beads with 2x 80% EtOH washes, and the sample was eluted in 10 µl of elution buffer. Upon incubation overnight with rotation at 4 °C and washing steps, ChIP DNA was isolated and set up in sequential reactions of adaptor fill-in, in vitro transcription, RNA 3’ adapter ligation, reverse transcription and PCR amplification to generate final libraries for each ChIP. After quality assessment and concentration estimation, libraries were combined and sequenced on an MGI DNBSEQ-G400RS instrument platform with paired-end settings.

##### Data Analysis

Raw ChIP-seq data was converted to FASTQ format and demultiplexed using mgikit (v. 0.1.4; settings: -m 1), allowing for up to one mismatch in the barcodes. Each FASTQ file of the demultiplexed samples were concatenated across the four lanes, creating the sample-specific FASTQ files for further processing. Quality evaluation, mapping, scaling the data to input, and the creation of bigWig files was done using the quantitative workflow (v. 0.6.0; settings: fragment_size: 400, max_barcode_errors: 1, mapping_quality: 0). The mapping was done using the GRCg6a reference genome, and low mapping regions were excluded. The scaled and pooled sample bigWig files for the modifications H3K27ac, H3K4me3, H3K9ac and H4K16ac, as well as the merged and low mapping regions excluded bigWig files from the ATAC, were ran through ChromHMM-tools to create signal input for use in ChromHMM. The signal input obtained from ChromHMM-tools was binarized and HMM models were created with ChromHMM (v. 1.25) using six states, which was mapped to galgal6.

#### Metaphase spreads and karyotyping of CEF samples

Metaphase spreads and karyotyping was performed as previously described (*60*), with slight modifications. Briefly, CEFs were grown as described above to a confluency of 70% and treated with CEF media containing 0.1µg/ml colcemid [Gibco] for 80 min at 37°C, 5% CO2. After 80 min, the treatment was removed, and the cells were washed with HBSS [Gibco] and detached using TryPLE [Gibco]. The cell suspension was then collected in media and centrifuged at 200g for 10 min at room temperature. The supernatant was removed, leaving 0.5ml in which the cell pellet was gently resuspended. For cell swelling, 10 ml of freshly prepared pre-warmed 0.075M KCl solution was added to the resuspended cells dropwise and the samples were incubated at 37°C for 12 min with gentle agitation every 2 min. The samples were then centrifuged at 200g for 5 min at room temperature. The supernatant was removed and discarded leaving 0.5 ml in which the cell pellet was gently resuspended. To fix the cells, 5 ml of freshly prepared Carnoy’s fixative (3:1 ratio methanol:acetic acid [Sigma]) was added while vortexing, followed by an additional 5ml added without vortexing. The samples were centrifuged at 200g for 5 min at room temperature. The above-mentioned step was repeated by adding 5 ml of Carnoy’s fixative without vortexing. For the final fixation, the supernatant was discarded, leaving 0.5ml in which the pellet was gently resuspended and 5 ml of Carnoy’s fixative was added, and the samples were stored at +4°C until slide preparation. *Slide preparation*. Microscope slides were first placed in a Coplin jar filled with absolute ethanol [VWR] for 10 min and rinsed in distilled water to prepare them for metaphase spreading. The cells were resuspended in freshly prepared Carnoy’s fixative and dropped onto the prepared microscope slides from a distance of 5cm. The slide was fixed with a large drop of Carnoy’s fixative and left to dry at room temperature. Freshly prepared Giemsa solution (3:1 ratio of Gurr buffer [Gibco]: Giemsa stain [Gibco]) was added to the slides for 10 min at room temperature. The slides were rinsed in distilled water and left to dry at room temperature. Coverslips [Brand] were then mounted using Permount [Sigma] and slides observed under 40x or 100x brightfield microscope objectives.

## References

1. J. A. M. Graves, Evolution of vertebrate sex chromosomes and dosage compensation. Nat Rev Genet 17, 33–46 (2016).

2. H. J. Muller, THE RELATION OF RECOMBINATION TO MUTATIONAL ADVANCE. Mutat Res 106, 2–9 (1964).

3. J. Felsenstein, The evolutionary advantage of recombination. Genetics 78, 737–756 (1974).

4. S. Ohno, Sex Chromosomes and Sex-linked Genes. In Monographs on endocrinology. Springer-Verlag, Heidelberg-Berlin-New York 1 (1967).

5. P. R. Baverstock, M. Adams, R. W. Polkinghorne, M. Gelder, A sex-linked enzyme in birds--Z-chromosome conservation but no dosage compensation. Nature 296, 763–766 (1982).

6. H. Ellegren, L. Hultin-Rosenberg, B. Brunström, L. Dencker, K. Kultima, B. Scholz, Faced with inequality: chicken do not have a general dosage compensation of sex-linked genes. BMC Biol 5, 40 (2007).

7. E. Melamed, A. P. Arnold, Regional differences in dosage compensation on the chicken Z chromosome. Genome Biol 8, R202 (2007).

8. J. E. Mank, H. Ellegren, All dosage compensation is local: gene-by-gene regulation of sex-biased expression on the chicken Z chromosome. Heredity (Edinb) 102, 312–320 (2009).

9. J. B. Wolf, J. Bryk, General lack of global dosage compensation in ZZ/ZW systems? Broadening the perspective with RNA-seq. BMC Genomics 12, 91 (2011).

10. S. Uebbing, A. Konzer, L. Xu, N. Backström, B. Brunström, J. Bergquist, H. Ellegren, Quantitative Mass Spectrometry Reveals Partial Translational Regulation for Dosage Compensation in Chicken. Mol Biol Evol 32, 2716–2725 (2015).

11. M. Teranishi, Y. Shimada, T. Hori, O. Nakabayashi, T. Kikuchi, T. Macleod, R. Pym, B. Sheldon, I. Solovei, H. Macgregor, S. Mizuno, Transcripts of the MHM region on the chicken Z chromosome accumulate as non-coding RNA in the nucleus of female cells adjacent to the DMRT1 locus. Chromosome Research 9, 147–165 (2001).

12. M. Warnefors, K. Mössinger, J. Halbert, T. Studer, J. L. VandeBerg, I. Lindgren, A. Fallahshahroudi, P. Jensen, H. Kaessmann, Sex-biased microRNA expression in mammals and birds reveals underlying regulatory mechanisms and a role in dosage compensation. Genome Res. 27, 1961–1973 (2017).

13. Y. Cheng, Z. Zhang, G. Zhang, L. Chen, C. Zeng, X. Liu, Y. Feng, The Male-Biased Expression of miR-2954 Is Involved in the Male Pathway of Chicken Sex Differentiation. Cells 12, 4 (2022).

14. A. Fallahshahroudi, L. Rodríguez-Montes, S. Y. Taemeh, N. Trost, M. Tellez, M. Ballantyne, A. Idoko-Akoh, L. Taylor, A. Sherman, E. Sorato, M. Johnsson, M. C. Moreira, M. J. McGrew, H. Kaessmann, A male-essential microRNA is key for avian sex chromosome dosage compensation. bioRxiv, 2024.03.06.581755 (2024).

15. A. Lentini, B. Reinius, Limitations of X:autosome ratio as a measurement of X-chromosome upregulation. Current Biology 33, R395–R396 (2023).

16. Y. Itoh, E. Melamed, X. Yang, K. Kampf, S. Wang, N. Yehya, A. Van Nas, K. Replogle, M. R. Band, D. F. Clayton, E. E. Schadt, A. J. Lusis, A. P. Arnold, Dosage compensation is less effective in birds than in mammals. J Biol 6, 2 (2007).

17. P. P. Singh, J. Arora, H. Isambert, Identification of Ohnolog Genes Originating from Whole Genome Duplication in Early Vertebrates, Based on Synteny Comparison across Multiple Genomes. PLoS Comput Biol 11, e1004394 (2015).

18. F. Zimmer, P. W. Harrison, C. Dessimoz, J. E. Mank, Compensation of Dosage-Sensitive Genes on the Chicken Z Chromosome. Genome Biol Evol 8, 1233–1242 (2016).

19. D. W. Bellott, H. Skaletsky, T.-J. Cho, L. Brown, D. Locke, N. Chen, S. Galkina, T. Pyntikova, N. Koutseva, T. Graves, C. remitzki, W. C. Warren, A. G. Clark, E. Gaginskaya, R. K. Wilson, D. C. Page, Avian W and mammalian Y chromosomes convergently retained dosage-sensitive regulators. Nat Genet 49, 387–394 (2017).

20. Q. Zhou, J. Zhang, D. Bachtrog, N. An, Q. Huang, E. D. Jarvis, M. T. P. Gilbert, G. Zhang, Complex evolutionary trajectories of sex chromosomes across bird taxa. Science 346, 1246338 (2014).

21. D. K. Nguyen, C. M. Disteche, Dosage compensation of the active X chromosome in mammals. Nat. Genet. 38, 47–53 (2006).

22. H. Lin, J. A. Halsall, P. Antczak, L. P. O’Neill, F. Falciani, B. M Turner, Relative overexpression of X-linked genes in mouse embryonic stem cells is consistent with Ohno’s hypothesis. Nature Genetics 43, 1169–1170 (2011).

23. A. J. M. Larsson, C. Coucoravas, R. Sandberg, B. Reinius, X-chromosome upregulation is driven by increased burst frequency. Nat Struct Mol Biol 26, 963–969 (2019).

24. A. Lentini, H. Cheng, J. C. Noble, N. Papanicolaou, C. Coucoravas, N. Andrews, Q. Deng, M. Enge, B. Reinius, Elastic dosage compensation by X-chromosome upregulation. Nat Commun 13, 1854 (2022).

25. M. Thorne, R. Collins, B. Sheldon, Triploidy and other chromosomal abnormalities in a selected line of chickens. Genet Sel Evol 23, S212 (1991).

26. P. Stenberg, J. Larsson, Buffering and the evolution of chromosome-wide gene regulation. Chromosoma 120, 213–225 (2011).

27. M. Lin, M. H. Thorne, I. C. Martin, B. L. Sheldon, R. C. Jones, Development of the gonads in the triploid (ZZW and ZZZ) fowl, Gallus domesticus, and comparison with normal diploid males (ZZ) and females (ZW). Reprod Fertil Dev 7, 1185–1197 (1995).

28. B. Kumar, S. J. Elsässer, Quantitative Multiplexed ChIP Reveals Global Alterations that Shape Promoter Bivalency in Ground State Embryonic Stem Cells. Cell Rep 28, 3274–3284.e5 (2019).

29. M. Hagemann-Jensen, C. Ziegenhain, P. Chen, D. Ramsköld, G.-J. Hendriks, A. J. M. Larsson, O. R. Faridani, R. Sandberg, Single-cell RNA counting at allele and isoform resolution using Smart-seq3. Nat Biotechnol 38, 708–714 (2020).

30. B. Reinius, J. E. Mold, D. Ramsköld, Q. Deng, P. Johnsson, J. Michaëlsson, J. Frisén, R. Sandberg, Analysis of allelic expression patterns in clonal somatic cells by single-cell RNA-seq. Nat Genet 48, 1430–1435 (2016).

31. A. J. M. Larsson, C. Ziegenhain, M. Hagemann-Jensen, B. Reinius, T. Jacob, T. Dalessandri, G.-J. Hendriks, M. Kasper, R. Sandberg, Transcriptional bursts explain autosomal random monoallelic expression and affect allelic imbalance. PLoS Comput Biol 17, e1008772 (2021).

32. M. Hagemann-Jensen, C. Ziegenhain, R. Sandberg, Scalable single-cell RNA sequencing from full transcripts with Smart-seq3xpress. Nat Biotechnol 40, 1452–1457 (2022).

33. C. Ziegenhain, G.-J. Hendriks, M. Hagemann-Jensen, R. Sandberg, Molecular spikes: a gold standard for single-cell RNA counting. Nat Methods 19, 560–566 (2022).

34. J. Kim, J. C. Marioni, Inferring the kinetics of stochastic gene expression from single-cell RNA-sequencing data. Genome Biol 14, R7 (2013).

35. B. Reinius, R. Sandberg, Random monoallelic expression of autosomal genes: stochastic transcription and allele-level regulation. Nat. Rev. Genet. 16, 653–664 (2015).

36. A. J. M. Larsson, P. Johnsson, M. Hagemann-Jensen, L. Hartmanis, O. R. Faridani, B. Reinius, \AAsa Segerstolpe, C. M. Rivera, B. Ren, R. Sandberg, Genomic encoding of transcriptional burst kinetics. Nature 565, 251–254 (2019).

37. R. D. Dar, B. S. Razooky, A. Singh, T. V. Trimeloni, J. M. McCollum, C. D. Cox, M. L. Simpson, L. S. Weinberger, Transcriptional burst frequency and burst size are equally modulated across the human genome. Proc Natl Acad Sci U S A 109, 17454–17459 (2012).

38. D. Yean, J. Gralla, Transcription Reinitiation Rate: a Special Role for the TATA Box. Molecular and Cellular Biology 17, 3809–3816 (1997).

39. K. Tantale, F. Mueller, A. Kozulic-Pirher, A. Lesne, J.-M. Victor, M.-C. Robert, S. Capozi, R. Chouaib, V. Bäcker, J. Mateos-Langerak, X. Darzacq, C. Zimmer, E. Basyuk, E. Bertrand, A single-molecule view of transcription reveals convoys of RNA polymerases and multi-scale bursting. Nat Commun 7, 12248 (2016).

40. D. Nicolas, B. Zoller, D. M. Suter, F. Naef, Modulation of transcriptional burst frequency by histone acetylation. Proc Natl Acad Sci U S A 115, 7153–7158 (2018).

41. N. S. Fechheimer, Origins of heteroploidy in chicken embryos. Poult Sci 60, 1365–1371 (1981).

42. J. a. M. Graves, Sex and death in birds: a model of dosage compensation that predicts lethality of sex chromosome aneuploids. Cytogenet Genome Res 101, 278–282 (2003).

43. M.-L. Faucillion, J. Larsson, Increased expression of X-linked genes in mammals is associated with a higher stability of transcripts and an increased ribosome density. Genome Biol Evol 7, 1039–1052 (2015).

44. Z.-Y. Wang, E. Leushkin, A. Liechti, S. Ovchinnikova, K. Mößinger, T. Brüning, C. Rummel, F. Grützner, M. Cardoso-Moreira, P. Janich, D. Gatfield, B. Diagouraga, B. de Massy, M. E. Gill, A. H. F. M. Peters, S. Anders, H. Kaessmann, Transcriptome and translatome co-evolution in mammals. Nature 588, 642–647 (2020).

45. E. Yildirim, R. I. Sadreyev, S. F. Pinter, J. T. Lee, X-chromosome hyperactivation in mammals via nonlinear relationships between chromatin states and transcription. Nature Structural and Molecular Biology 19, 56–62 (2012).

46. X. Deng, J. B. Berletch, W. Ma, D. K. Nguyen, J. B. Hiatt, W. S. Noble, J. Shendure, C. M. Disteche, Mammalian X upregulation is associated with enhanced transcription initiation, RNA half-life, and MOF-mediated H4K16 acetylation. Developmental Cell 25, 55–68 (2013).

47. I. Talon, A. Janiszewski, B. Theeuwes, T. Lefevre, J. Song, G. Bervoets, L. Vanheer, N. De Geest, S. Poovathingal, R. Allsop, J.-C. Marine, F. Rambow, T. Voet, V. Pasque, Enhanced chromatin accessibility contributes to X chromosome dosage compensation in mammals. Genome Biol 22, 302 (2021).

48. N. C. Lister, A. M. Milton, H. R. Patel, S. A. Waters, B. J. Hanrahan, K. L. McIntyre, A. M. Livernois, W. B. Horspool, L. K. Wee, A. R. Ringel, S. Mundlos, M. I. Robson, L. Shearwin-Whyatt, F. Grützner, J. A. M. Graves, A. Ruiz errera, P. D. Waters, Incomplete transcriptional dosage compensation of chicken and platypus sex chromosomes is balanced by post-transcriptional compensation. Proc Natl Acad Sci U S A 121, e2322360121 (2024).

49. S. Parekh, C. Ziegenhain, B. Vieth, W. Enard, I. Hellmann, zUMIs - A fast and flexible pipeline to process RNA sequencing data with UMIs. GigaScience 7 (2018).

50. G. van der Auwera, B. D. O’Connor, Genomics in the Cloud: Using Docker, GATK, and WDL in Terra (O’Reilly Media, Sebastopol, CA, First edition., 2020).

51. P. Danecek, J. K. Bonfield, J. Liddle, J. Marshall, V. Ohan, M. O. Pollard, A. Whitwham, T. Keane, S. A. McCarthy, R. M. Davi, H. Li, Twelve years of SAMtools and BCFtools. GigaScience 10, giab008 (2021).

52. G. de Sena Brandine, A. D. Smith, Fast and memory-efficient mapping of short bisulfite sequencing reads using a two-letter alphabet. NAR Genom Bioinform 3, lqab115 (2021).

53. Q. Song, B. Decato, E. E. Hong, M. Zhou, F. Fang, J. Qu, T. Garvin, M. Kessler, J. Zhou, A. D. Smith, A Reference Methylome Database and Analysis Pipeline to Facilitate Integrative and Comparative Epigenomics. PLoS ONE 8, e81148 (2013).

54. C. Pockrandt, M. Alzamel, C. S. Iliopoulos, K. Reinert, GenMap: ultra-fast computation of genome mappability. Bioinformatics 36, 3687–3692 (2020).

55. S. Picelli, A. K. Björklund, B. Reinius, S. Sagasser, G. Winberg, R. Sandberg, Tn5 transposase and tagmentation procedures for massively scaled sequencing projects. Genome Res. 24, 2033–2040 (2014).

56. S. Chen, Y. Zhou, Y. Chen, J. Gu, fastp: an ultra-fast all-in-one FASTQ preprocessor. Bioinformatics 34, i884–i890 (2018).

57. H. Li, Minimap2: pairwise alignment for nucleotide sequences. Bioinformatics 34, 3094–3100 (2018).

58. A. R. Quinlan, I. M. Hall, BEDTools: a flexible suite of utilities for comparing genomic features. Bioinformatics 26, 841–842 (2010).

59. M. R. Corces, A. E. Trevino, E. G. Hamilton, P. G. Greenside, N. A. Sinnott-Armstrong, S. Vesuna, A. T. Satpathy, A. J. Rubin, K. S. Montine, B. Wu, A. Kathiria, S. W. Cho, M. R. Mumbach, A. C. Carter, M. Kasowski, L. A. Orloff, V. I. Risca, A. Kundaje, P. A. Khavari, T. J. Montine, W. J. Greenleaf, H. Y. Chang, An improved ATAC-seq protocol reduces background and enables interrogation of frozen tissues. Nat Methods 14, 959–962 (2017).

60. B. Howe, A. Umrigar, F. Tsien, Chromosome preparation from cultured cells. J Vis Exp, e50203 (2014).

